# Positioning of nucleosomes containing γ-H2AX precedes active DNA demethylation and transcription initiation

**DOI:** 10.1101/2020.03.06.980912

**Authors:** Stephanie Dobersch, Karla Rubio, Indrabahadur Singh, Stefan Günther, Johannes Graumann, Julio Cordero, Rafael Castillo-Negrete, Minh Bao Huynh, Aditi Mehta, Peter Braubach, Hector Cabrera-Fuentes, Jürgen Bernhagen, Cho-Ming Chao, Saverio Bellusci, Andreas Günther, Klaus T Preissner, Gergana Dobreva, Malgorzata Wygrecka, Thomas Braun, Dulce Papy-Garcia, Guillermo Barreto

## Abstract

In addition to nucleosomes, chromatin contains non-histone chromatin-associated proteins, of which the high-mobility group (HMG) proteins are the most abundant. Chromatin-mediated regulation of transcription involves DNA methylation and histone modifications. However, the order of events and the precise function of HMG proteins during transcription initiation remain unclear. Here we show that HMG AT-hook 2 protein (HMGA2) induces DNA nicks at the transcription start site, which are required by the histone chaperone FACT (facilitates chromatin transcription) complex to incorporate nucleosomes containing the histone variant H2A.X. Further, phosphorylation of H2A.X at S139 (γ-H2AX) is required for repair-mediated DNA demethylation and transcription activation. The relevance of these findings is demonstrated within the context of TGFB1 signaling and idiopathic pulmonary fibrosis, suggesting therapies against this lethal disease. Our data support that chromatin opening during transcriptional initiation involves intermediates with DNA breaks that subsequently require DNA repair mechanisms to ensure the integrity of the genome.

## INTRODUCTION

In the eukaryotic cell nucleus, chromatin is the physiological template of all DNA-dependent processes including transcription. The structural and functional units of chromatin are the nucleosomes, each one consisting of ∼147 bp of genomic DNA wrapped around a core histone octamer, which in turn is built of two H2A–H2B dimers and one (H3–H4)_2_ tetramer (Hall et al., 2009; Ozturk et al., 2014). In addition to canonical histones (H1, H2A, H2B, H3 and H4), there are so called histone variants for all histones except for H4. Histone variants differ from the canonical histones in their amino acid sequence and have specific and fundamental functions that cannot be performed by canonical histones. Histone variants are strongly conserved among species, suggesting that they arose at an early stage in evolution. The canonical histone H2A has a large number of variants, each with defined biochemical and functional properties (Bonisch and Hake, 2012; Gaume and Torres-Padilla, 2015). Here we focus on the histone variant H2AFX (commonly known as H2AX, further referred to as H2A.X), which represents about 2-25% of the cellular H2A pool in mammals (Rogakou et al., 1998). Phosphorylated H2A.X at serine 139 (H2A.XS139ph; commonly known as γ-H2AX, further referred to as pH2A.X) is used as a marker for DNA double-strand breaks (Redon et al., 2011). However, accumulating evidence suggests additional functions of pH2A.X (Singh et al., 2015; Steinel et al., 2013; Turinetto et al., 2012; Ziegler-Birling et al., 2009). The histone chaperone FACT (facilitates chromatin transcription) is a heterodimeric complex, consisting of SUPT16 and SSRP1 (Spt16 and Pob3 in yeast) that is responsible for the deposition of H2A/H2B-dimers onto DNA (Belotserkovskaya et al., 2003; Mason and Struhl, 2003). The FACT complex mainly interacts with H2B mediating the deposition of H2A/H2B-dimers containing different H2A variants (Hondele et al., 2013). Thus, the deposition of H2A.X into chromatin seems to be mediated by the FACT complex (Winkler and Luger, 2011).

In addition to nucleosomes, chromatin consists of non-histone chromatin associated proteins, of which the high-mobility group (HMG) proteins are the most abundant. Although HMG proteins do not possess intrinsic transcriptional activity, they are called architectural transcription factors because they modulate the transcription of their target genes by altering the chromatin structure at the promoter and/or enhancers (Reeves, 2010). HMG proteins are divided into three families based on their DNA binding domains: HMGA (containing AT-hooks), HMGB (containing HMG-boxes) and HMGN (containing nucleosomal binding domains) (Bustin, 2001; Ozturk et al., 2014). Here we will focus on HMG AT-hook 2 protein (HMGA2), a member of the HMGA family that mediates transforming growth factor beta 1 (TGFB1, commonly known as TGFβ1) signaling (Thuault et al., 2006). We have previously shown that HMGA2-induced transcription requires phosphorylation of H2A.X at S139, which in turn is mediated by the protein kinase ataxia telangiectasia mutated (ATM) (Singh et al., 2015). Furthermore, we demonstrated the biological relevance of this mechanism of transcriptional initiation within the context of TGFB1 signaling and epithelial-mesenchymal transition (EMT). Interestingly, TGFB signaling has been reported to induce active DNA demethylation with the involvement of thymidine DNA glycosylase (TDG) (Thillainadesan et al., 2012). Active DNA demethylation also requires GADD45A (growth arrest and DNA damage protein 45 alpha) and TET1 (ten-eleven translocation methylcytosine dioxygenase 1), which sequentially oxidize 5-methylcytosine (5mC) to 5-carboxylcytosine (5caC) (Arab et al., 2019; Barreto et al., 2007) and are cleared through DNA repair mechanisms. TGFB1 signaling and EMT are both playing a crucial role in idiopathic pulmonary fibrosis (IPF). IPF is the most common interstitial lung disease showing a prevalence of 20 new cases per 100,000 persons per year (Coward et al., 2010; Noble et al., 2012). A central event in IPF is the abnormal proliferation and migration of fibroblasts in the alveolar compartment in response to lung injury. Under normal circumstances fibroblasts are important for wound healing and connective tissue production. However, in the fibrotic lung their function is impaired resulting in formation of fibroblastic foci, which consist of highly proliferative fibroblasts, immune cells, and excessive extracellular matrix (ECM) protein deposition, such as fibronectin (FN1) and collagen (COL1A1) (Barkauskas and Noble, 2014). Consequently, these processes result in disproportionate levels of scar tissue, alterations of the lung epithelium structure and loss of the gas exchange function of the lung. IPF patients die within 2 years after diagnosis mostly due to respiratory failure. Current treatments against IPF aim to ameliorate patient symptoms and to delay disease progression (Selvaggio and Noble, 2016). Unfortunately, therapies targeting the causes of or reverting IPF have not yet been developed. Here we demonstrate that inhibition of the HMGA2-FACT-ATM-pH2A.X axis reduce fibrotic hallmarks *in vitro* using primary human lung fibroblast (hLF) and *ex vivo* using human precision-cut lung slices (hPCLS), both from control and IPF patients. Our study supports the development of therapeutic approaches against IPF using FACT inhibition.

## RESULTS

### HMGA2 is required for pH2A.X deposition at transcription start sites

We have previously reported that HMGA2-mediated transcription requires phosphorylation of the histone variant H2A.X at S139, which in turn is catalyzed by the protein kinase ATM (Singh et al., 2015). To further dissect this mechanism of transcription initiation, we decided first to determine the effect of *Hmga2*-knockout (KO) on genome wide levels of pH2A.X. We performed next generation sequencing (NGS) after chromatin immunoprecipitation (ChIP-seq; Figures 1 and S1A) using pH2A.X-specific antibodies and chromatin isolated from mouse embryonic fibroblasts (MEF) from wild-type (WT = *Hmga2*+/+) and *Hmga2*-deficient (*Hmga2*-KO = *Hmga2*−/−) embryos. The analysis of these ChIP-seq results using the UCSC Known Genes dataset (Hsu et al., 2006) revealed that pH2A.X is specifically enriched at transcription start sites (TSS) of genes in an *Hmga2*-dependent manner (Figure 1A), seeing as *Hmga2*-KO significantly reduced pH2A.X levels. A zoom into the −750 to +750 base pair (bp) region relative to the TSS (Figure 1B) revealed that pH2A.X levels significantly peaked at the TSS (−250 to +250 bp) in *Hmga2*+/+ MEF. Further, the genes were ranked based on pH2A.X levels at the TSS (Table S1) and the results were visualized as heat maps (Figure 1C). From the top 15% of the genes with high pH2A.X levels at TSS (further referred as top 15% candidates), we selected *Gata6* (GATA binding protein 6), *Mtor* (mechanistic target of rapamycin kinase) and *Igf1* (insulin like growth factor 1) for further single gene analysis. Explanatory for these gene selection, we have previously reported *Gata6* as direct target gene of HMGA2 (Singh et al., 2014; Singh et al., 2015), KEGG (Kyoto Encyclopedia of Genes and Genomes) pathway enrichment based analysis of the top 15% candidates showed significant enrichment of genes related to the mammalian target of rapamycin (mTOR) signaling pathway (Figure S1B) and HMGA2 has been related to the insulin signaling pathway (Brants et al., 2004; Li et al., 2012). Visualization of the selected genes using the UCSC genome browser confirmed the reduction of pH2A.X at specific regions close to TSS in *Hmga2-/-* MEF when compared to *Hmga2+/+* MEF (Figure 1D, top). Similar results were obtained after ChIP-seq using H2A.X and H3 antibodies (Figure 1D, bottom). Promoter analysis of *Gata6*, *Mtor* and *Igf1* by ChIP using pH2A.X-, H2A.X-, H3- and HMGA2-specific antibodies (Figure S1C-D) confirmed the ChIP-seq data. These findings suggest that the first nucleosome relative to the TSS of the top 15% candidates contains pH2A.X and *Hmga2* is required for correct positioning of this first nucleosome.

**Figure 1:**
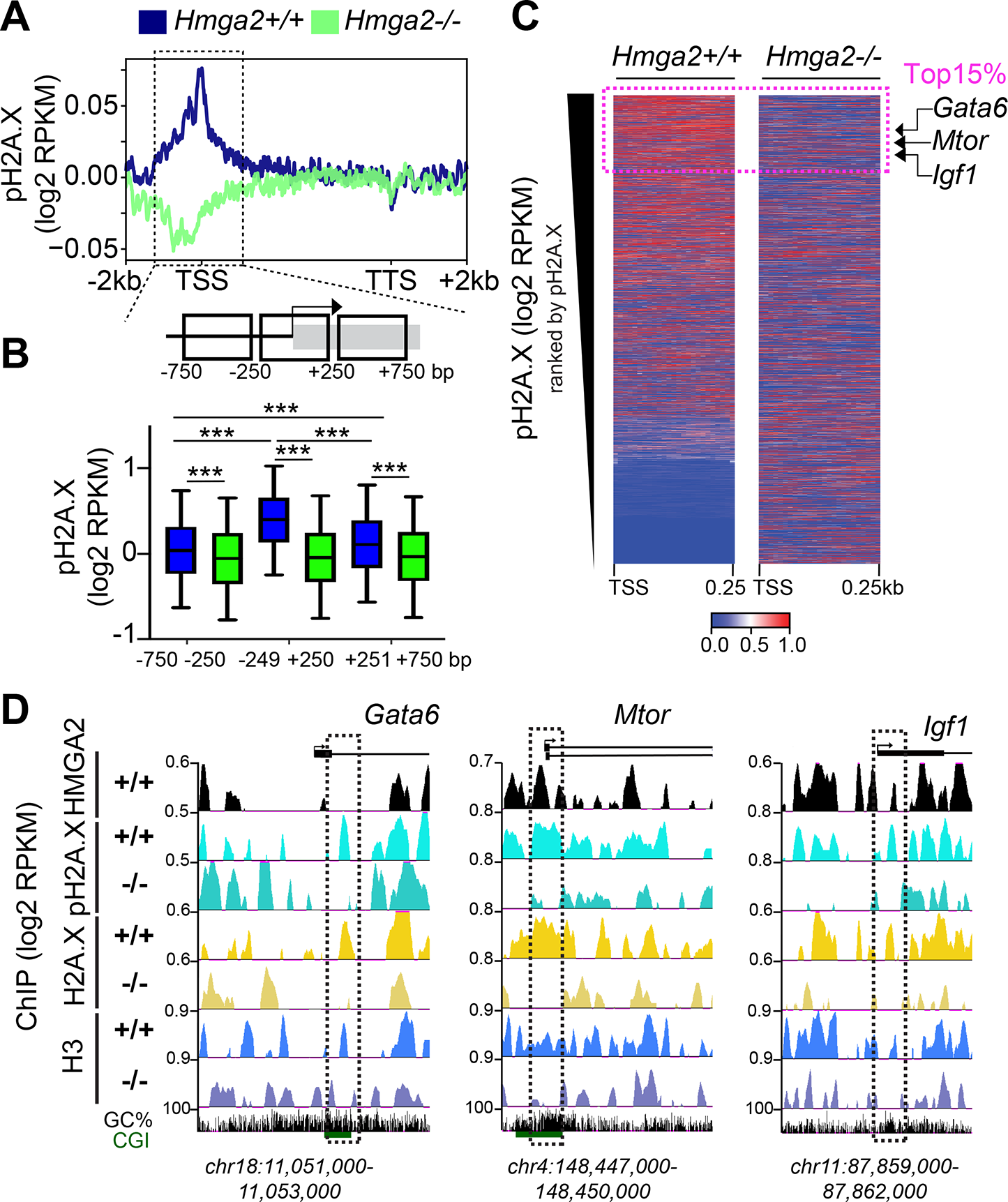
HMGA2 is required for pH2A.X deposition at TSS. (A) Aggregate plot for pH2A.X enrichment within the gene body ±2kb of UCSC known genes in *Hmga2+/+* and *Hmga2-/-* MEF. ChIP-seq reads were normalized using reads per kilobase per million (RPKM) measure and are represented as log2 enrichment over their corresponding inputs. TSS, transcription start site; TTS, transcription termination site. Dotted square, ±750bp region around the TSS. (B) Top, schematic representation of the genomic region highlighted in A. Bottom, box plot of pH2A.X enrichment in the genomic regions showed as squares at the top in *Hmga2+/+* and *Hmga2-/-* MEF. RPKM of the pH2A.X ChIP-seq were binned within each of these genomic regions and represented as log2. *P* values after Wilcoxon-Mann-Whitney test, *** *P* ≤ 0.001. The statistical test values are shown in Supplementary Table S3. (C) Heat map for pH2A.X enrichment at the TSS +0.25kb of UCSC known genes in *Hmga2+/+* and *Hmga2-/*-MEF. Genes were ranked by pH2A.X enrichment in *Hmga2+/+* MEF. Doted square, the top 15% ranked genes, as well as *Gata6*, *Mtor* and *Igf1* were selected for further analysis. (D) Visualization of selected HMGA2 target genes using UCSC Genome Browser showing HMGA2, pH2A.X, H2A.X and H3 enrichment in *Hmga2+/+* and -/- MEF. ChIP-seq reads were normalized using RPKM measure and are represented as log2 enrichment over their corresponding inputs. Images show the indicated gene loci with their genomic coordinates. Arrows, direction of the genes; black boxes, exons; dotted squares, regions selected for single gene analysis. See also Figure S1 and Table S1.

### Position of the first nucleosome containing pH2A.X determines the basal transcription activity of genes

Phosphorylation of specific amino acids in the C-terminal domain of the large subunit of the RNA polymerase II (Pol II) determines its interaction with specific factors, thereby regulating the transcription cycle consisting of initiation, elongation and termination (Kim et al., 2010). To monitor transcription initiation, ChIP-seq was performed using antibodies specific for transcription initiating S5 phosphorylated Pol II (further referred to as pPol II) and chromatin isolated from *Hmga2*+/+ and *Hmga2*−/− MEF (Figures 2 and S1A). Analysis of the ChIP-seq results using the UCSC Known Genes dataset revealed that pPol II was enriched at TSS in *Hmga2*-dependent manner (Figure 2A). In addition, we observed that pPol II enrichment coincides with pH2A.X peaks at TSS (Figure 2B) also in an *Hmga2*-dependent manner. Visualization of *Gata6*, *Mtor* and *Igf1* using the UCSC genome browser (Figure 2C) and ChIP analysis of their promoters (Figure 2D, left) confirmed the reduction of pPol II at specific regions close to TSS in *Hmga2-/-* MEF when compared to *Hmga2+/+* MEF. Furthermore, the reduced pPol II levels after *Hmga2*-KO correlated with the reduced expression of the analyzed genes as shown by quantitative reverse-transcription PCR (qRT-PCR, Figure 2D, right). Interestingly, analysis of the ChIP-seq data by *k*-means clustering (doi: 10.2307/2346830) revealed three clusters in the top 15% candidates (Figure 2E). Cluster 1 showed pPol II, pH2A.X and HMGA2 enrichment directly at the TSS (top), while clusters 2 and 3 showed enrichment of these proteins 125 bp and 250 bp 3′ of the TSS, respectively (middle and bottom). Further, RNA sequencing (RNA-seq) based expression analysis in *Hmga2*+/+ and *Hmga2-/-* MEF (Figure 2F) revealed that the genes in the three clusters have different basal transcription activities, whereby cluster 1 has the lowest, cluster 2 the middle and cluster 3 the highest basal transcription activity in *Hmga2+/+* MEF. *Hmga2*-KO significantly reduced the basal transcription activity in all three clusters. Our results support a correlation between the basal transcription activity and the position of pPol II, pH2A.X and HMGA2 relative to TSS.

**Figure 2:**
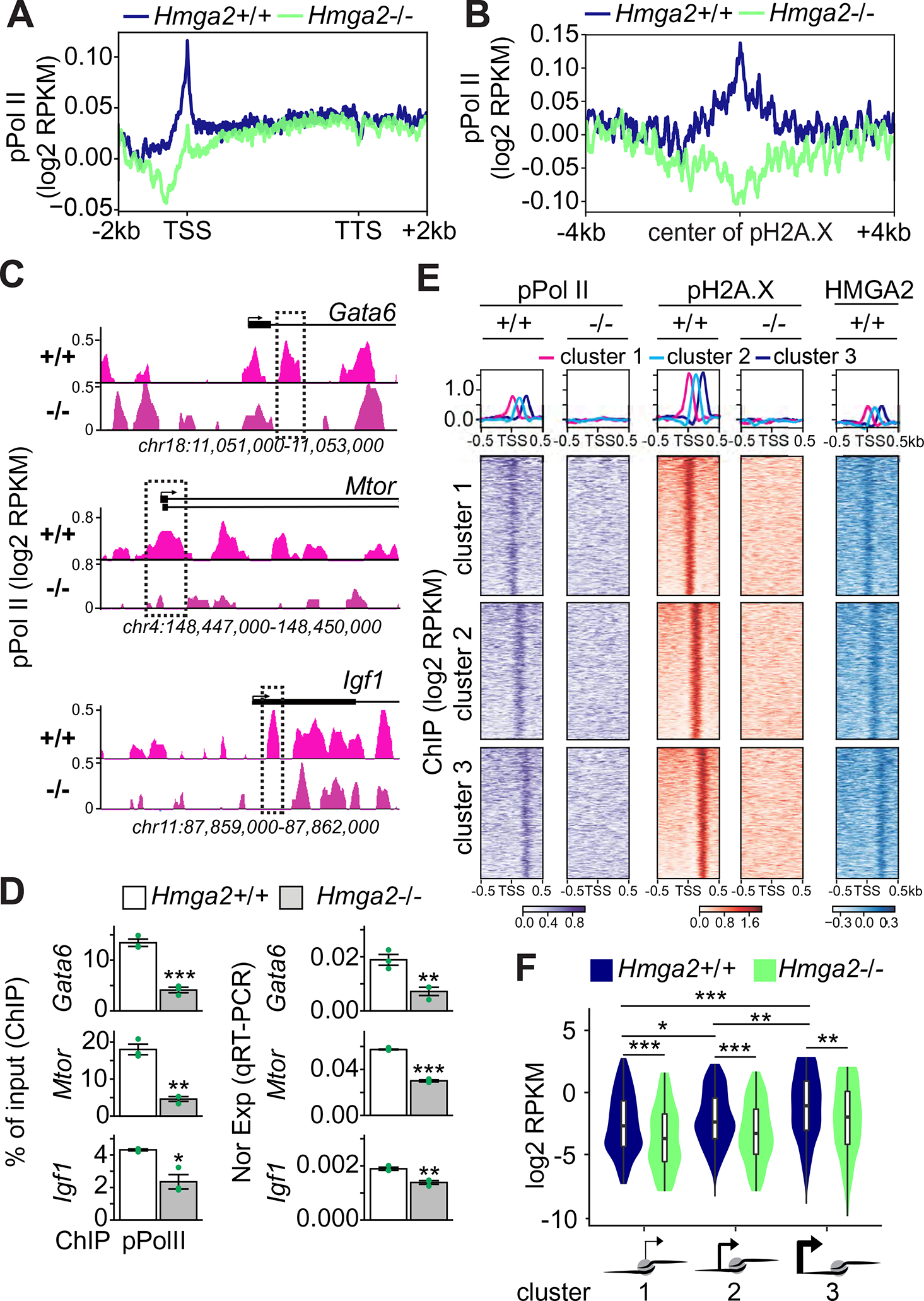
Position of first nucleosome containing pH2A.X determines basal transcription activity. (A-B) Aggregate plots for phosphorylated serine 5 RNA polymerase II (pPol II) enrichment within the gene body ±2kb of UCSC known genes (A) and in a ±4kb region respective to pH2A.X peaks (B) in *Hmga2+/+* and *Hmga2-/-* MEF. ChIP-seq reads were normalized using reads per kilobase per million (RPKM) measure and are represented as log2 enrichment over their corresponding inputs. TSS, transcription start site; TTS, transcription termination site. (C) Visualization of selected HMGA2 target genes using UCSC Genome Browser showing pPol II enrichment in *Hmga2+/+* and -/- MEF. ChIP-seq reads were normalized using RPKM measure and are represented as log2 enrichment over their corresponding inputs. Images represent the indicated gene loci with their genomic coordinates. Arrows, direction of the genes; black boxes, exons; dotted squares, regions selected for single gene analysis. (D) Analysis of selected HMGA2 target genes. Left, ChIP of *Gata6*, *Mtor* and *Igf1* after pPol II immunoprecipitation in *Hmga2+/+* and *Hmga2-/-* MEF. Right, qRT-PCR-based, *Tuba1a*-normalized expression analysis under the same conditions. Bar plots presenting data as means; error bars, s.e.m (*n* = 3 independent experiments); asterisks, *P* values after t-Test, *** *P* ≤ 0.001; ** *P* ≤ 0.01; * *P* ≤ 0.05. (E) Aggregate plots (top) and heat maps (bottom) for pPol II, pH2A.X and HMGA2 enrichment at the TSS ± 0.5kb of the top 15% candidates in *Hmga2+/+* and *Hmga2-/-* MEF. Three clusters were generated using *k*-means algorithm. Genes were sorted based on the enrichment of pH2A.X in *Hmga2+/+* MEF. (F) Violin and box plot representing the basal transcription activity (as log2 RPKM) of genes in *Hmga2+/+* and *Hmga2-/-* MEF. The genes were sorted in three groups based on the position of the first pH2A.X-containing nucleosome 5’ to the TSS. Asterisks, *P* values after Wilcoxon-Mann-Whitney test, *** *P* ≤ 0.001; ** *P* ≤ 0.01; * *P* ≤ 0.05. See also Figure S1 and Table S1. The statistical test values of each plot are shown in Supplementary Table S3.

### HMGA2 is required for enrichment of the FACT complex at TSS

To gain new insights in the HMGA2, ATM, and pH2A.X transcriptional network (Singh et al., 2015), native chromatin preparations from *Hmga2*+/+ and *Hmga2-/-* MEF were digested with micrococcal nuclease (MNase) and subsequently fractionated by sucrose gradient ultracentrifugation (SGU) (Figures 3A-E and S2A-B). From the obtained fractions, the proteins were extracted and analyzed either by western blot (WB; Figures 3A and 3D, top), or by high-resolution mass spectrometry-based proteomic approach (Figures 3B-C and Table S1), while the DNA was also isolated and analyzed either by gel electrophoresis (Figure 3D, bottom), or by DNA sequencing (MNase-seq, Figure 3E and S2C-D). In *Hmga2*+/+ MEF (Figure 3A, left), WB of the obtained fractions showed that HMGA2 sedimented in fractions 4 to 9, whereas pPol II and histones mainly sedimented in fractions 1 to 4, where protein complexes of higher molecular weight (MW) are expected. Interestingly, H2A.X showed a similar sedimentation pattern as the core histones H2A, H2B and H3, whereas pH2A.X mainly sedimented in fraction 4 together with HMGA2. On the other hand, in *Hmga2*-/- MEF (Figure 3A, right) were only minor variations in the sedimentation pattern of all tested proteins, with the most notable change being the reduction in the levels of HMGA2 and pH2A.X compared to *Hmga2*+/+ MEF. The subsequent analysis was focused on fractions 3 and 4, because fraction 4 contained all proteins monitored by WB in *Hmga2*+/+ MEF, whereas fraction 3 contained all proteins tested by WB besides HMGA2 and pH2A.X. We analyzed these two fractions by high-resolution mass spectrometry-based proteomic approach and identified proteins that were more than 1.65-fold significantly enriched in *Hmga2*+/+ MEF when compared to *Hmga2*-/- MEF (Figure 3B; Table S1; *n*=1,215 and *P*˂0.05 in fraction 3; *n*=1,729 and *P*˂0.05 in fraction 4). A closer look on the proteins enriched in fractions 3 and 4 of *Hmga2+/+* MEF revealed the presence of both components of the FACT complex, SUPT16 and SSRP1 (Figure 3B), as well as proteins related to transcription regulation and nucleotide excision repair (NER; Figure S2B). Interestingly, *Hmga2*-KO significantly reduced the levels of SUPT16 and SSRP1 in fraction 4 without significantly affecting their levels in fraction 3 (Figure 3C). These results were confirmed by WB of SGU fractions using SUPT16- or SSRP1-specific antibodies (Figure 3D, top). Further, we isolated and analyzed by electrophoresis the DNA from the SGU fractions and found DNA fragments with a length profile of 100-300 bp that mainly sedimented in fractions 3 to 5 of both *Hmga2*+/+ and *Hmga2-/-* MEF. The length profile of DNA fragments supports the majority presence of mono- and di-nucleosomes in the native chromatin preparations that were fractionated by SGU. Focusing again on fraction 3 and 4, MNase-seq was performed and reads between 100 to 200 bp were selected for downstream analysis (Figures 3E and S2C-D). Using the UCSC Known Genes as reference dataset, we found that in fraction 4 the sequencing reads were enriched with TSS in an *Hmga2*-dependent manner (Figure 3E), since this enrichment was abolished after *Hmga2*-KO. In fraction 3, we did not detect TSS enrichment of the sequencing reads. Our results indicate that the native chromatin in fraction 4 contains mono- and di-nucleosomes (Figure S2D), which are enriched with TSS, HMGA2, pH2A.X and pPoll II in WT MEF. To link the results obtained by MNase-seq and mass spectrometry after fractionation by SGU, we performed ChIP-seq using SUPT16- and SSRP1-specific antibodies and chromatin from *Hmga2*+/+ and *Hmga2-/-* MEF (Figures 3F and S2E). Confirming the results in fraction 4, we detected accumulation of both components of the FACT complex at TSS in *Hmga2+/+* MEF, whereas *Hmga2*-KO reduced the levels of SUPT16 and SSRP1 at TSS. In summary, our results demonstrated that HMGA2 is required for binding of the FACT complex to TSS.

**Figure 3:**
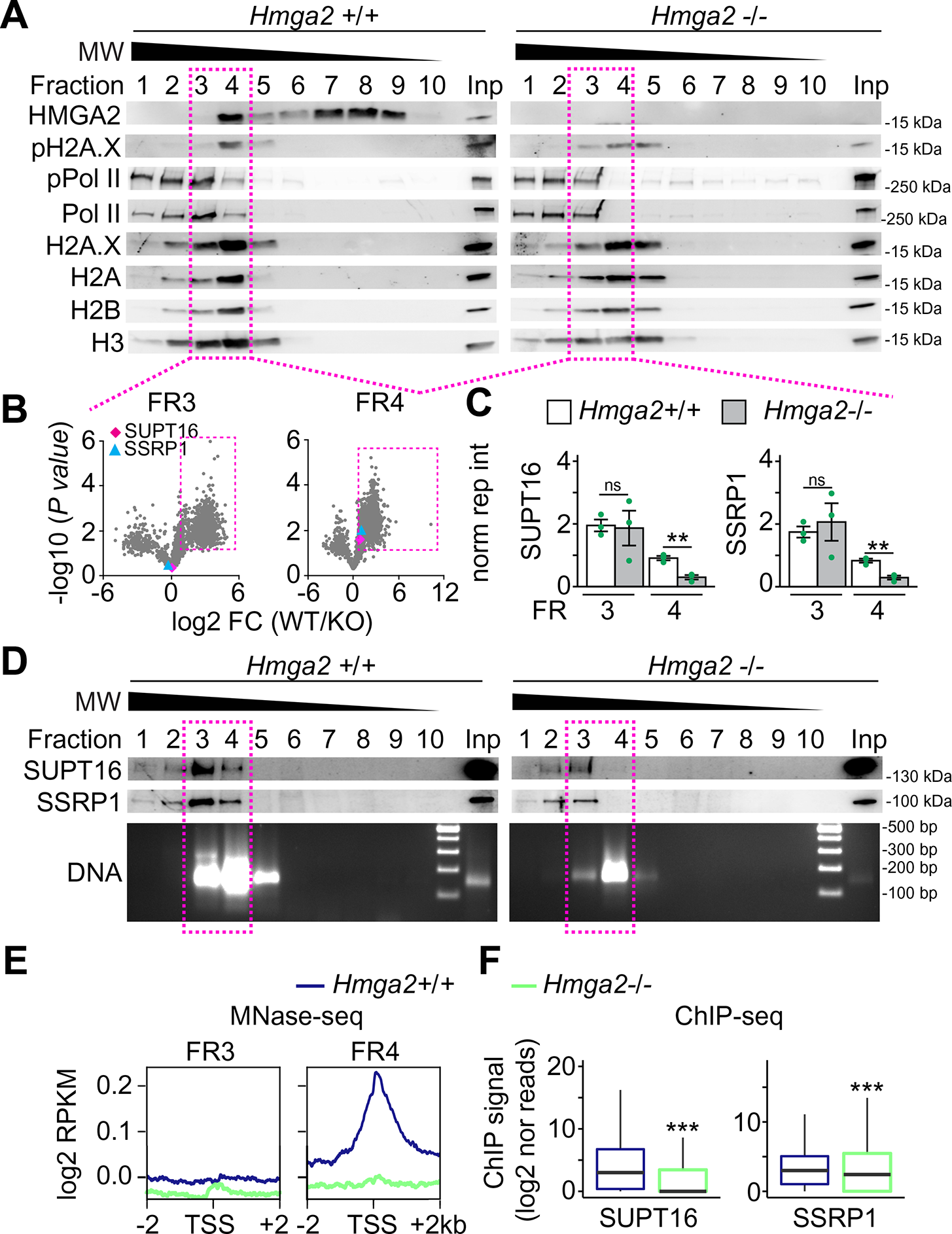
HMGA2 is required for enrichment of the FACT complex at TSS. (A-E) Native chromatin from *Hmga2+/*+ and -/- MEF was digested with micrococcal nuclease (MNase) and fractionated by sucrose gradient ultracentrifugation (SGU). (A) The obtained fractions were analyzed by WB using the indicated antibodies. MW, molecular weight, kDa, kilo Dakton. Inp, input represents 0.5% of the material used for SGU. Square, fractions selected for further analysis. (B) Mass spectrometry analysis of proteins in fractions 3 and 4. Volcano plot representing the significance (-log10 *P*-values after one-tailed *t*-test) vs. intensity fold change between *Hmga2+/+* and *-/-* MEF (log2 of means intensity ratios from three independent experiments). Square, proteins with log2 fold change ≥1. Diamond, SUPT16; triangle, SSRP1. (C) Bar plots showing normalized reporter intensity of SUPT16 (left) and SSRP1 (right) in fractions 3 and 4 of the SGU in A. Data are shown as means ± s.e.m. (*n* = 3 independent experiments); asterisks, *P* values after *t*-Test, ** *P* ≤ 0.01; ns, non-significant. (D) Top, WB analysis as in A using antibodies specific for components of the FACT complex. Bottom, DNA was isolated from the fractions obtained by SGU in A and analyzed by agarose gel electrophoresis. Square, fractions selected for MNase-seq. (E) MNase-seq of fractions 3 and 4 of the SGU in A. Aggregate plots representing the enrichment over input (as log2 RPKM) of genomic sequences relative to the TSS ±2kb. (F) Box plots of ChIP-seq-based SUPT16 (left) and SSRP1 (right) enrichment analysis within the TSS + 0.5kb of the top 15% candidates in *Hmga2+/+* and *-/-* MEF. Values are represented as log2 of mapped reads that were normalized to the total counts and the input was subtracted. Boxes, interquartile ranges; whiskers, 5 to 95% confidence intervals; horizontal lines, medians; asterisks, *P* values after two-tailed Mann-Whitney test, *** *P* ≤ 0.001; ns, not significant. See also Figure S2. The statistical test values of each plot are shown in Supplementary Table S3.

### HMGA2-FACT interaction and -lyase activity are required for pH2A.X deposition and solving of R-loops

The results in Figures 3A-F suggest an interaction between HMGA2 and the FACT complex. In addition, we found in our previously published mass spectrometry based HMGA2 interactome (Singh et al., 2015) that HMGA2 precipitated SUPT16 and SSRP1 (Figure S2F). Indeed, the interaction of HMGA2 with both components of the FACT complex was confirmed by co-immunoprecipitation (Co-IP) assay using nuclear protein extracts from MEF after overexpression of HMGA2 tagged C-terminally with MYC and HIS (HMGA2-MYC-HIS; Figure 4A). To further characterize the HMGA2-FACT interaction, nuclear extracts from *Hmga2*+/+ and *Hmga2-/-* MEF were fractionated into chromatin-bound and nucleoplasm fractions (Figure S3A). WB of these fractions showed that HMGA2 was exclusively bound to chromatin, whereas SUPT16 and SSRP1 were present in both sub-nuclear fractions. However, higher levels of both FACT components were detected in the chromatin-bound fraction of *Hmga2*+/+ MEF as compared to the nucleoplasm. Interestingly, *Hmga2*-KO reverted the distribution of SUPT16 and SSRP1 between these two sub-nuclear fractions, thereby supporting that *Hmga2* is required for tethering the FACT complex to chromatin. These results were confirmed by ChIP of *Gata6*, *Mtor* and *Igf1* (Figure 4B) using SUPT16- and SSRP1-specific antibodies and chromatin isolated from *Hmga2*+/+ and *Hmga2*-/- MEF that were stably transfected with a tetracycline-inducible expression construct (tetOn) either empty (-; negative control) or containing the cDNA of WT HMGA2-MYC-HIS. Doxycycline treatment of these stably transfected MEF induced the expression of WT HMGA2-MYC-HIS (Figure S4B). *Hmga2*-KO reduced the levels of SUPT16 and SSRP1 in all promoters analyzed (Figure 4B), while doxycycline-inducible expression of WT HMGA2-MYC-HIS in *Hmga2*-/- MEF reconstituted the levels of both FACT components, thereby demonstrating the specificity of the effects caused by *Hmga2*-KO. To demonstrate the causal involvement of the FACT complex in context of HMGA2-mediated chromatin rearrangements, we analyzed the levels of pH2A.X and H2A.X at the *Gata6*, *Mtor* and *Igf1* promoters by ChIP using chromatin from *Hmga2*+/+ MEF and stably transfected *Hmga2*-/- MEF that were treated with DMSO (control) or a FACT inhibitor (FACTin; CBLC000 trifluoroacetate) (Chang et al., 2018) and doxycycline as indicated (Figure 4C). FACTin treatment induced chromatin trapping of SUPT16 and SSRP1 (Figure S3B-C) (Gasparian et al., 2011). In addition, FACT inhibition significantly reduced pH2A.X and H2A.X levels at the analyzed promoters, confirming that the FACT complex is required for proper H2A.X deposition (Figure 4C, left). Further, *Hmga2*-KO also reduced pH2A.X and H2A.X levels at the analyzed promoters (middle), while doxycycline-inducible expression of WT HMGA2-MYC-HIS in *Hmga2*-/- MEF reconstituted the pH2A.X and H2A.X levels (right), thereby confirming the results presented in Figures 1 and 4B. Remarkably, FACT inhibition counteracted the reconstituting effect mediated by doxycycline-inducible expression of WT HMGA2-MYC-HIS, showing the causal involvement of the FACT complex in the function of HMGA2. In summary, our results demonstrated that the FACT complex is required for HMGA2 function and consequently also for proper pH2A.X levels at the analyzed promoters.

**Figure 4:**
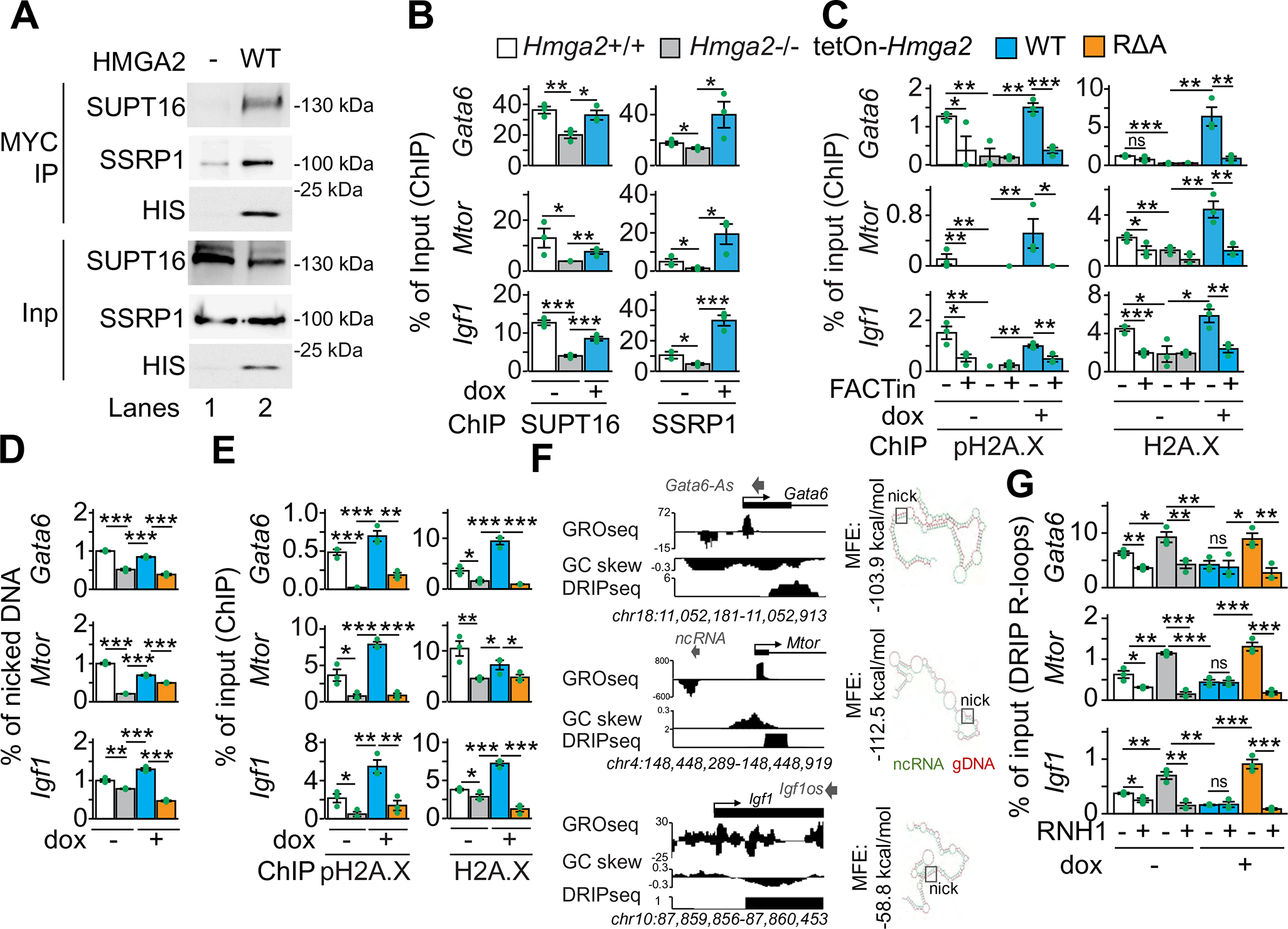
HMGA2-FACT interaction and -lyase activity are required for pH2A.X deposition and solving of R-loops. (A) Western blot using the indicated antibodies after co-immunoprecipitation (Co-IP) assay using nuclear protein extracts from *Hmga2-/-* MEF that were non-transfected (-) or stably transfected with *Hmga2-myc-his* (WT) and magnetic beads coated with MYC-specific antibodies. Input, 5% of IP starting material. (B-C, E) ChIP-based promoter analysis of selected HMGA2 target genes using the indicated antibodies and chromatin from *Hmga2+/+*, *Hmga2-/-* MEF, as well as *Hmga2-/-* MEF that were stably transfected with a tetracycline-inducible expression construct (tetOn) either for WT *Hmga2-myc-his* or the lyase-deficient mutant RΔA *Hmga2-myc-his*. MEF were treated with doxycycline and FACT inhibitor (FACTin; CBLC000 trifluoroacetate) as indicated. (D) Analysis of single-strand DNA breaks at promoters of selected HMGA2 target genes using genomic DNA from MEF as in E. (F) Left, Genome-browser visualization of selected HMGA2 target genes showed nascent RNA (GRO-seq) in WT MEF (Busslinger et al., 2017), GC-skew and DNA-RNA hybrids (DRIP-seq) in NIH/3T3 mouse fibroblasts (Sanz et al., 2016). Images represent mapped sequence tag densities relative to the indicated loci. Genomic coordinates are shown at the bottom. Arrow heads, non-coding RNAs in antisense orientation; Arrows, direction of the genes; black boxes, exons. Right, *in silico* analysis revealed complementary sequences between the identified antisense ncRNA (green) and genomic sequences at the TSS of the corresponding mRNAs (red) with relatively favorable minimum free energy (MFE), supporting the formation of DNA-RNA hybrids containing a nucleotide sequence that favors DNA nicks (squares). (G) Analysis of selected HMGA2 target genes by DNA–RNA immunoprecipitation (DRIP) using the antibody S9.6 (Boguslawski et al., 1986) and nucleic acids isolated from MEF treated as in E. Prior IP, nucleic acids were digested with RNase H1 (RNH1) as indicated. In all bar plots, data are shown as means ± s.e.m. (*n* = 3 independent experiments); asterisks, *P* values after *t*-Test, *** *P* ≤ 0.001; ** *P* ≤ 0.01; * *P* ≤ 0.05; ns, non-significant. See also Figures S3 and S4. The statistical test values of each plot are shown in Supplementary Table S3.

It has been reported that HMGA2 efficiently cleaves DNA generating single-strand breaks (Summer et al., 2009). The arginine residues of the C-terminal AT-hook motif are crucial for this intrinsic lyase activity of HMGA2 (Figure S4A, top). Thus, to further elucidate the molecular mechanism of HMGA2-mediated transcription activation, a lyase-deficient HMGA2 mutant was generated by substituting these arginine residues by alanine (RΔA HMGA2; Figure S4A, bottom). Further, *Hmga2*-/- MEF were stably transfected with a tetracycline-inducible expression construct containing the cDNA of RΔA HMGA2-MYC-HIS. Doxycycline treatment of these stably transfected MEF induced the expression of RΔA HMGA2-MYC-HIS to similar levels as in MEF containing WT HMGA2-MYC-HIS (Figure S4B, top). Furthermore, analysis of the chromatin-bound sub-nuclear fraction of these cells (Figure S4B, bottom) demonstrated that the R to A mutations did not significantly affect the binding of HMGA2 to chromatin. In addition, the loss of lyase activity in RΔA HMGA2-MYC-HIS was confirmed by monitoring DNA damage using Comet Assays (Jachimowicz et al., 2019) (Figure S4C) or by monitoring HMGA2-DNA complexes on dot blots (Figure S4D) after trapping experiments (Summer et al., 2009) using NaCNBH_3_ as reducing agent to trap the Schiff base intermediates of the lyase reaction mediated by HMGA2. Furthermore, to demonstrate that the R to A mutations in the third AT-hook domain of HMGA2 do not affect the interaction with the FACT complex, we performed Co-IP of nuclear protein extracts from the stably transfected *Hmga2*-/- MEF described above (Figure S4E). IP using nuclear extracts and MYC-specific antibodies specifically co-precipitated both components of the FACT complex, SUPT16 and SSRP1, without significant differences between WT and RΔA HMGA2-MYC-HIS. Moreover, we analyzed the promoters of *Gata6*, *Mtor* and *Igf1* (Figure S4F) by ChIP using chromatin isolated from the same cells used for the Co-IP (Figure S4E). We found that the enrichment of WT and RΔA HMGA2-MYC-HIS was not significantly different at the *Igf1* promoter, whereas we observed an increased enrichment of RΔA HMGA2-MYC-HIS at the *Gata6* and *Mtor* promoters when compared to WT HMGA2-MYC-HIS. These results indicate that the interaction with the FACT complex and the enrichment at the analyzed promoters was similar for WT and RΔA HMGA2.

After confirming the loss of lyase activity in RΔA HMGA2-MYC-HIS (Figures S4C-D), single-strand DNA breaks (DNA nicks) at the *Gata6*, *Mtor* and *Igf1* promoters were monitored using genomic DNA from *Hmga2*+/+ MEF and stably transfected *Hmga2*-/- MEF that were non-treated (-) or treated with doxycycline (Figure 4D). We detected DNA nicks at the analyzed promoters in *Hmga2*+/+ MEF, whose levels were reduced upon *Hmga2*-KO. Interestingly, inducible expression of WT HMGA2 in *Hmga2-/-* MEF reconstituted the levels of DNA nicks, whereas RΔA HMGA2 did not rescue the effect induced by *Hmga2*-KO, thereby confirming that HMGA2 lyase activity is required for the DNA nicks detected at the analyzed promoters. Further, we decided to demonstrate the requirement of the lyase activity for the function of HMGA2. Thus, the levels of pH2A.X and H2A.X at the *Gata6*, *Mtor* and *Igf1* promoters were analyzed by ChIP using chromatin from *Hmga2*+/+ and stably transfected *Hmga2*-/- MEF (Figure 4E). Confirming the results in Figures 1 and 4C, *Hmga2*-KO reduced pH2A.X and H2A.X levels at the analyzed promoters. In addition, doxycycline-inducible expression of WT HMGA2 in *Hmga2*-/- cells reconstituted the levels of pH2A.X and H2A.X, demonstrating the specificity of the effect observed after *Hmga2*-KO. However, doxycycline-inducible expression of RΔA HMGA2 in *Hmga2*-/- MEF did not rescue the effect induced by *Hmga2*-KO. Although the mutations inducing the loss of the lyase activity did not affect the HMGA2-FACT complex interaction (Figure S4E), these results showed that the lyase activity is required for proper pH2A.X and H2A.X levels at the analyzed promoters. Interestingly, genome-wide run-on assay (GRO-seq) in WT MEF (Busslinger et al., 2017) revealed nascent noncoding RNAs (ncRNAs) in antisense orientation from 46% of the loci (*n*= 4,380) of the top 15% candidates (*n* = 9,522; Figures S4H-I), including *Gata6*, *Mtor* and *Igf1* (Figure 4F, left top). *In silico* analysis allowed us to detect putative binding sites of the identified ncRNAs at the TSS of the corresponding mRNAs with relatively favorable minimum free energy (< − 55 kcal/mol; Figure 4F, right), supporting the formation of DNA-RNA hybrids containing a nucleotide sequence that favors DNA nicks (Gutjahr and Xu, 2014). In the same genomic regions, we also identified strand asymmetry in the distribution of cytosines and guanines, so called GC skews (Figure 4F, left middle; Figure S4H, bottom), that are predisposed to form R-loops, which are three-stranded nucleic acid structures consisting of a DNA-RNA hybrid and the associated non-template single-stranded DNA (Skourti-Stathaki and Proudfoot, 2014). Supporting this hypothesis, published genome-wide sequencing experiments after DNA–RNA immunoprecipitation (DRIP-seq) in NIH/3T3 mouse fibroblasts (Sanz et al., 2016) confirmed the formation of DNA-RNA hybrids in 38% of the promoters (*n* = 3,618) of the top 15% candidates (*n* = 9,522; Figures S4H-I), including *Gata6*, *Mtor* and *Igf1* (Figure 4F, left bottom). All these observations prompted us to investigate the role of HMGA2 during R-loop formation at the TSS. Thus, we analyzed by DRIP assays the levels of R-loops at the *Gata6*, *Mtor* and *Igf1* promoters using the antibody S9.6 (Boguslawski et al., 1986) and nucleic acids isolated from *Hmga2*+/+ and stably transfected *Hmga2*-/- MEF (Figure 4G). *Hmga2*-KO increased R-loops levels at the promoters analyzed, whereas doxycycline-inducible expression of WT HMGA2 in *Hmga2*-/- MEF reduced R-loop levels back to similar levels as in *Hmga2*+/+ MEF. Interestingly, doxycycline-inducible expression of RΔA HMGA2 in *Hmga2*-/- MEF did not rescue the effect induced by *Hmga2*-KO. In parallel, treatment of the samples before IP with RNase H1 (RNH1), which degrades RNA in DNA-RNA hybrids, reduced the levels of R-loops in all tested conditions, demonstrating the specificity of the antibody S9.6 (Boguslawski et al., 1986). In summary, our results demonstrated that HMGA2 and its lyase activity are required to solve R-loops at the analyzed promoters.

### HMGA2-FACT-ATM-pH2A.X axis is required to solve R-loops and induce DNA demethylation

The inducible expression of RΔA HMGA2 in *Hmga2*-/- MEF did not decrease R-loops levels at TSS that were increased after *Hmga2*-KO (Figure 4G), supporting that the lyase activity of HMGA2 is required to solve R-loops. To further investigate these results, the levels of double-stranded DNA (dsDNA) at the *Gata6*, *Mtor* and *Igf1* promoters were analyzed by DNA immunoprecipitation (DIP) assays (Figure 5A, left). Inversely correlating with the effects on R-loops, *Hmga2*-KO reduced dsDNA levels at the promoters analyzed. Further, doxycycline-inducible expression of WT HMGA2 in *Hmga2-/-* MEF reconstituted dsDNA levels, whereas RΔA HMGA2 failed to rescue the effect induced by *Hmga2*-KO. These results further support the requirement of HMGA2 and its lyase activity for solving R-loops. Since DNA methylation alters chromatin structure and is associated with R-loop formation (Arab et al., 2019; Black and Whetstine, 2011), we also analyzed the levels of 5-methylcytosine (5mC) at the *Gata6*, *Mtor* and *Igf1* promoters by DIP assays using 5mC-specific antibodies (Mould et al., 2013) (Figure 5A, right). Correlating with the effects on R-loops, *Hmga2*-KO increased 5mC levels, which in turn were reduced by inducible expression of WT HMGA2 in *Hmga2*-/- cells but not by RΔA HMGA2. These results showed that HMGA2 and its lyase activity are required for proper 5mC levels at the analyzed promoters. In addition, the results using FACTin (Figure 4C) showed that the FACT complex is required for HMGA2 function and consequently for proper pH2A.X levels at TSS. Thus, to demonstrate the sequential order of events of the molecular mechanism proposed here (Figure 5B), additional experiments were performed (Figures 5 and S5). We first analyzed the *Gata6*, *Mtor* and *Igf1* promoters by DRIP and DIP using nucleic acids isolated from *Hmga2*+/+ MEF that were non-treated (-) or treated with FACTin as indicated (Figures S5A-B). FACTin treatment in *Hmga2+/+* MEF increased R-loop and 5mC levels, whereas dsDNA levels were reduced, thereby supporting that the FACT complex is required to solve R-loops and for proper levels of 5mC at the analyzed promoters, similarly as the HMGA2 lyase activity (Figures 4G and 5A). Previously, we have shown that ATM loss-of-function (LOF) blocks TGFB1-induced and HMGA2-mediated transcription activation (Singh et al., 2015). To confirm the causal involvement of ATM in the mechanism of transcription regulation proposed here (Figure 5B), the levels of pH2A.X and H2A.X at the *Gata6*, *Mtor* and *Igf1* promoters were analyzed by ChIP using chromatin from *Hmga2*+/+ MEF and stably transfected *Hmga2*-/- MEF that were treated with DMSO (control) or an ATM inhibitor (ATMi; KU-55933) and doxycycline as indicated (Figure 5C). Interestingly, ATMi treatment counteracted the rescue effect on pH2A.X levels that was mediated by inducible expression of WT HMGA2 in *Hmga2-/-* MEF, without significantly affecting H2A.X levels, thereby supporting that ATM is required for the post-translational modification of H2A.X rather than for the deposition of H2A.X into the analyzed promoters. In addition, we monitored 5mC levels by DIP at the *Gata6*, *Mtor* and *Igf1* promoters (Figure 5D) and found that ATM-LOF also counteracted the rescue effect on 5mC levels mediated by inducible expression of WT HMGA2 in *Hmga2-/-* MEF, thereby supporting that phosphorylation of H2A.X at S139 is required for proper 5mC levels. The results obtained after ATM-LOF (Figures 5C-D) support that ATM acts downstream of HMGA2 and the FACT complex. To gain further insights into the order of events proposed here (Figure 5B), the effect of *Gadd45a*-specific LOF was analyzed using small interfering RNA (siRNA; *siG45a*; Figure S5C) on pH2A.X, H2A.X and 5mC levels at the *Gata6*, *Mtor* and *Igf1* promoters in *Hmga2*+/+ MEF and stably transfected *Hmga2*-/- MEF (Figures 5E-F). While *siG45a* transfection counteracted the rescue effect on 5mC levels mediated by inducible expression of WT HMGA2 in *Hmga2-/-* MEF (Figure 5F), it did not significantly affect pH2A.X and H2A.X levels (Figure 5E), confirming that GADD45A is required for proper 5mC levels but not for pH2A.X and H2A.X levels. Further, we found that GADD45A gain-of-function (GOF) after transfection of a human *GADD45A* expression construct into mouse lung epithelial (MLE-12) cells reduced 5mC levels in HMGA2-dependent manner (Figures S5D-E). Our results (Figures 5E-F and S5D-E) indicate that GADD45A acts downstream of the HMGA2-FACT-ATM-pH2A.X axis (Figure 5B). Confirming this interpretation, ChIP-seq using GADD45A-specific antibodies and chromatin isolated from *Hmga2+/+* and *Hmga2-/-* MEF (Figure S5F) revealed that GADD45A and pH2A.X are enriched at similar regions respective to TSS of the top 15% candidates. Moreover, ChIP analysis of the *Gata6*, *Mtor* and *Igf1* promoters using GADD45A- or TET1-specific antibodies and chromatin from *Hmga2*+/+ and stably transfected *Hmga2*-/- MEF that were treated with DMSO (control) or doxycycline (Figure 5G) showed that *Hmga2*-KO abrogated GADD45A and TET1 binding to the analyzed promoters. Strikingly, inducible expression of WT HMGA2 reconstituted GADD45A and TET1 binding to the analyzed promoters, whereas RΔA HMGA2 did not rescue the effect induced by *Hmga2*-KO.

**Figure 5:**
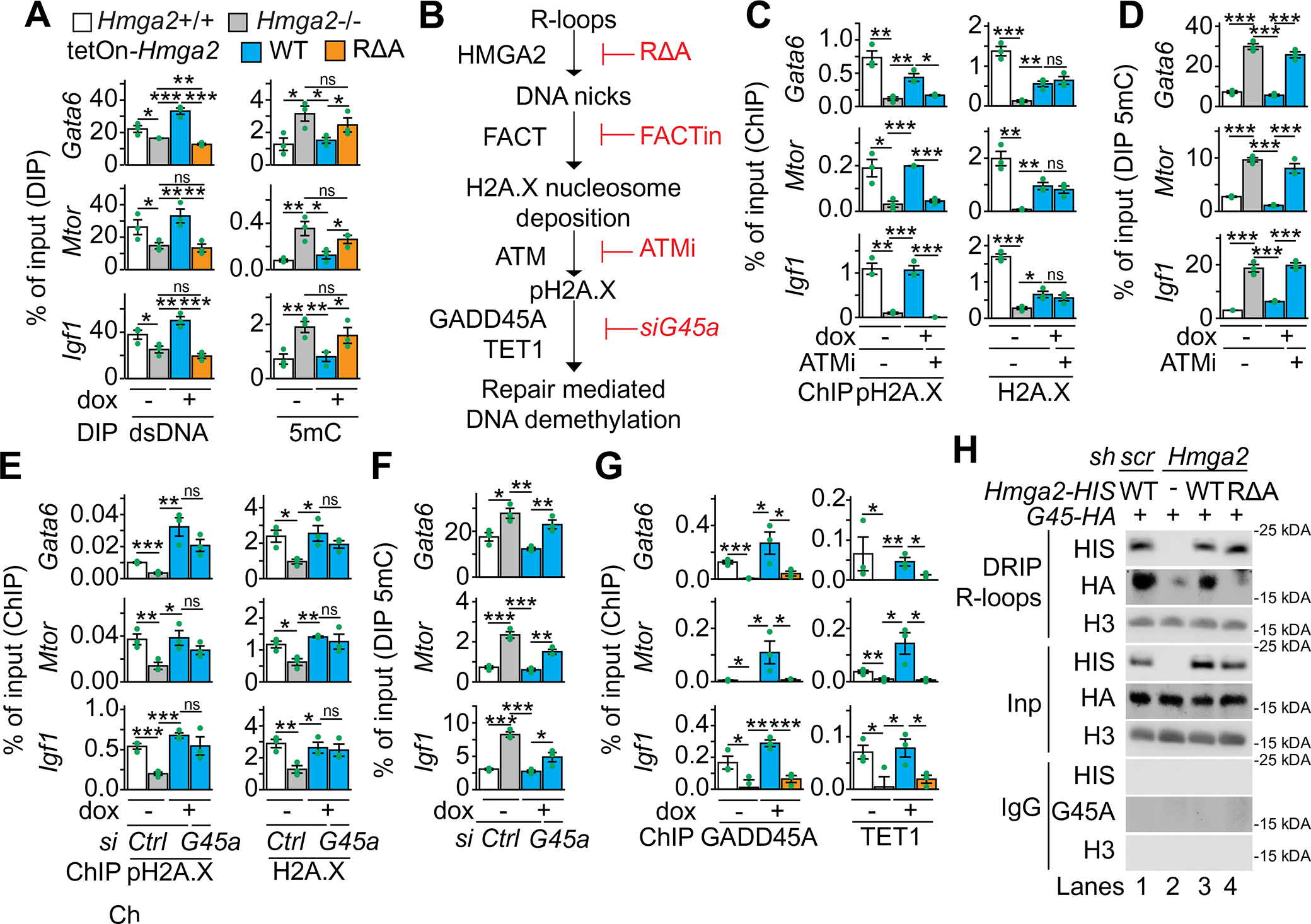
HMGA2-FACT-ATM-pH2A.X axis is required to solve R-loops and induce DNA demethylation. (A) DNA immunoprecipitation (DIP) based promoter analysis of selected HMGA2 target genes using antibodies specific for double-stranded DNA (dsDNA) or 5-methylcytosine (5mC) and genomic DNA from *Hmag2+/+*, *Hmag2-/-* MEF, as well as *Hmga2-/-* MEF that were stably transfected with a tetracycline-inducible expression construct (tetOn) for either WT *Hmga2-myc-his* or the lyase-deficient mutant RΔA *Hmga2-myc-his*. MEF were treated with doxycycline as indicated. (B) Schematic representation of the sequential order of events during transcription activation mediated by the HMGA2-FACT-ATM-pH2A.X axis. (C) ChIP-based promoter analysis of selected HMGA2 target genes using the indicated antibodies and chromatin from MEF treated as in A. In addition, MEF were treated with ATM inhibitor (ATMi; KU-55933) as indicated. (D) DIP-based promoter analysis as in A, using 5mC-specific antibodies. In addition, MEF were treated with ATMi as indicated. (E) ChIP-based promoter analysis of selected HMGA2 target genes using the indicated antibodies and chromatin from MEF treated as in A. In addition, MEF were transfected with control (Ctrl) or *Gadd45a*-specific small interfering RNA (siRNA) as indicated. (F) DIP-based promoter analysis as in A, using 5mC-specific antibodies. In addition, MEF were transfected with Ctrl or *Gadd45a*-specific siRNA as indicated. (G) ChIP-based promoter analysis of selected HMGA2 target genes using the indicated antibodies and chromatin from MEF treated as in A. (H) WB analysis using antibodies specific for HIS-tag, HA-tag and H3 after DRIP using the antibody S9.6 (Boguslawski et al., 1986) and chromatin isolated from MLE-12 cells that were stably transfected either with a control (scramble, *scr*) or an *Hmga2*-specific short hairpin DNA (*sh*) construct and transiently transfected with WT *Hmga2-myc-his* or the lyase-deficient mutant RΔA *Hmga2-myc-his* and *Gadd45*-HA as indicated. Input (Inp), 5% of IP starting material; immunoglobulin G (IgG), negative control. In all bar plots, data are shown as means ± s.e.m. (*n* = 3 independent experiments); asterisks, *P* values after *t*-Test, *** *P* ≤ 0.001; ** *P* ≤ 0.01; * *P* ≤ 0.05; ns, non-significant. See also Figure S5. The statistical test values of each plot are shown in Supplementary Table S3.

We have shown that genetic ablation of *Hmga2* increased R-loop levels (Figure 4G) and reduced GADD45A binding (Figure 5G) at the *Gata6*, *Mtor* and *Igf1* promoters. Interestingly, Arab and colleagues recently reported that GADD45A preferentially binds DNA-RNA hybrids and R-loops rather than single-stranded (ss) or double-stranded (ds) DNA or RNA (Arab et al., 2019). To elucidate these at first glance contradictory results, we performed DRIP using the antibody S9.6 (Boguslawski et al., 1986) and chromatin from stably transfected *Hmga2*-/- MEF that were non-treated (-) or treated with doxycycline to induce WT or RΔA HMGA2 (Figure 5H). WB analysis of the precipitated material revealed that both WT and RΔA HMGA2 bind to R-loops. Further, GADD45A also binds to R-loops, confirming the results by Arab and colleagues (Arab et al., 2019). However, GADD45A binding to R-loops increased after inducible expression of WT HMGA2, but not after RΔA HMGA2, suggesting that DNA nicks in the R-loops increase the affinity of GADD45A to the R-loops. Our results by WB after DRIP (Figure 5H) correlated with the ChIP analysis of the *Gata6*, *Mtor* and *Igf1* promoters (Figure 5G) and were confirmed by DRIP and sequential ChIP (DRIP-ChIP; Figure S5G). Taking together, our results demonstrate that the HMGA2-FACT-ATM-pH2A.X axis acts upstream of GADD45A and facilitates its binding to R-loops at specific promoters by nicking the DNA moiety of the DNA-RNA hybrid, thereby inducing DNA repair-mediated promoter demethylation.

### HMGA2-FACT-ATM-pH2A.X axis mediates TGFB1 induced transcription activation

We have previously shown that HMGA2 mediates TGFB1 induced transcription (Singh et al., 2015). Thus, we decided to evaluate the mechanism of transcription activation proposed here (Figure 5B) within the context of TGFB1 signaling. We performed RNA-seq in *Hmga2*+/+ and *Hmga2*−/− MEF that were non-treated or treated with TGFB1 and visualized the results of those genes that were induced by TGFB1 treatment as heat maps after *k*-means clustering (Figures 6A and S6A). Four clusters were identified, of which clusters 2 (*n* = 1,471) and 4 (*n* = 1,974) contained genes that were TGFB1 inducible in *Hmga2* independent manner. Cluster 3 (*n* = 381) contained TGFB1 inducible genes, whose expression increased after *Hmga2*-KO, while TGFB1 treatment in *Hmga2-/-* MEF reduced their expression. We focused on cluster 1 (*n* = 640) for further analysis, which contained TGFB1 inducible genes in *Hmga2* dependent manner. Cross-analysis of our RNA-seq after TGFB1 treatment (Figure 6A and S6A) with our ChIP-seq data (Figures 2E and S5F) confirmed the existence of three gene groups based on the position of the first nucleosome 3’ of the TSS containing pH2A.X, which we called position clusters 1 to 3 to differentiate them from the TGFB1 inducible clusters. Consistent with our previous results (Figure 2F), the genes in the position clusters 1 to 3 displayed increasing basal transcription activity from position cluster 1 with the lowest to position cluster 3 with the highest (Figure S6B). In addition, the position of the first nucleosome relative to the TSS also correlated with the strength of transcriptional activation induced by TGFB1 (Figure 6B), where position cluster 1 showed the lowest, cluster 2 a medium and cluster 3 the highest transcriptional inducibility by TGFB1 in *Hmga2+/+* MEF. Remarkably, *Hmga2*-KO reduced the inducibility of the genes after TGFB1 treatment in all three position clusters. ChIP-seq analysis of pH2A.X levels was also performed using the same conditions as in our RNA-seq after TGFB1 treatment (Figures 6C and S6C). TGFB1 treatment increased pH2A.X levels at the TSS of TGFB1 inducible cluster 1 genes in *Hmga2+/+* MEF, whereas this effect was not observed in *Hmga2-/-* MEF, confirming the requirement of *Hmga2* for the effects induced by TGFB1. Further, to determine the causal involvement of the FACT complex during TGFB1 induced transcriptional activation, we performed a series of experiments analyzing *Gata6*, *Mtor* and *Igf1* in *Hmga2+/+* and *Hmga2-/-* MEF that were non-treated or treated with FACTin (Figures 6D-F). TGFB1 treatment in *Hmga2+/+* MEF increased the expression of *Gata6*, *Mtor* and *Igf1* (Figure 6D) as well as the levels of pPol II and pH2A.X in their promoters (Figure 6E), whereas 5mC levels were reduced (Figure 6F). The effects induced by TGFB1 treatment were not observed in *Hmga2-/-* MEF confirming the requirement of *Hmga2*. Further, FACTin treatment counteracted the effects induced by TGFB1 in *Hmga2+/+* MEF supporting the causal involvement of the FACT complex. Interestingly, WB analysis of protein extracts from *Hmga2+/+* and *Hmga2-/-* MEF (Figure S6D) demonstrated that the effects observed after *Hmga2*- and FACT-LOF take place neither affecting total SMAD2/3 levels, nor their activation by TGFB1. Consistent with the mechanism of transcriptional regulation proposed here (Figure 5B) and the results in the Figures 5E-F, siRNA-mediated *Gadd45a*-LOF counteracted the reducing effect of TGFB1 on 5mC levels in *Hmga2+/+* MEF (Figure 6G) without affecting the increasing effect on pH2A.X levels (Figure 6H), thereby confirming that GADD45A acts downstream of the HMGA2-FACT-ATM-pH2A.X axis.

**Figure 6:**
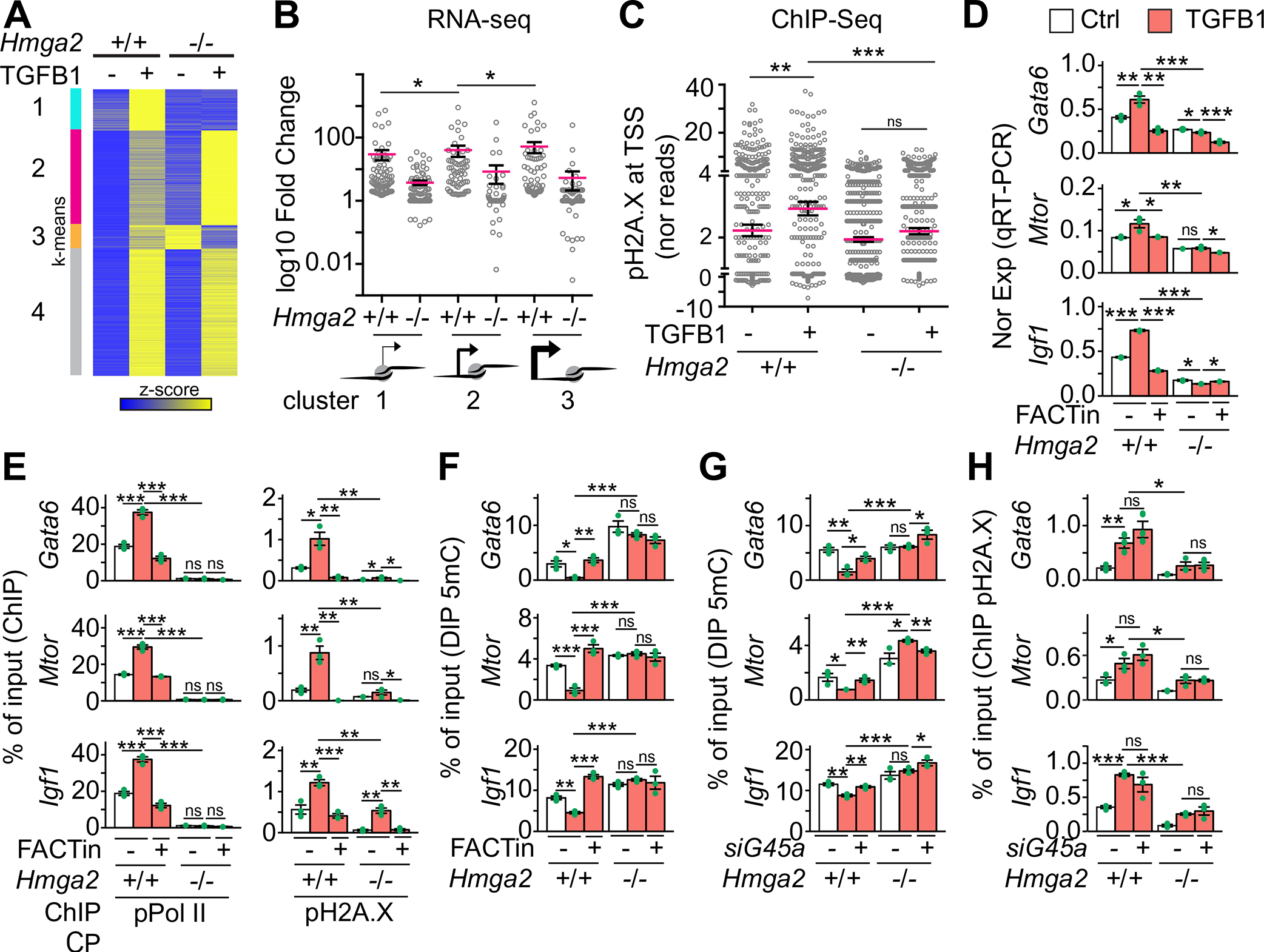
HMGA2-FACT-ATM-pH2A.X axis mediates TGFB1 induced transcription activation. (A) Heat map showing RNA-seq-based expression analysis of TGFB1-inducible genes in *Hmga2+/+* and *Hmga2-/-* MEF non-treated or treated with TGFB1. Data were normalized by Z score transformation and clustered using *k*-means algorithm. (B) Dot plot presenting the RNA-seq-based expression analysis of cluster 1 genes from A as log10 fold change between TGFB1-treated and non-treated MEF. Genes were grouped into the position clusters identified in Figure 2E. Each dot represents the value of a single gene. Red line, average; error bars, s.e.m.; asterisks, *P* values after unpaired Mann-Whitney Test, * *P* ≤ 0.05. (C) Dot plot presenting ChIP-seq-based pH2A.X enrichment analysis in the top 15% candidates from Figure 1C using chromatin from MEF treated as in A. For each gene, normalized reads in the TSS +0.25kb region were binned and the maximal value was plotted. Inputs were subtracted from the corresponding samples. Red line, average; error bars, s.e.m.; asterisks, *P* values after unpaired Mann-Whitney Test, *** *P* ≤ 0.001; ** *P* ≤ 0.01; ns, non-significant. (D) qRT-PCR-base expression analysis of HMGA2 target genes in *Hmag2+/+*, *Hmag2-/-* MEF that were non-treated (Ctrl) or treated with TGFB1 and FACT inhibitor (FACTin; CBLC000 trifluoroacetate) as indicated. (E) ChIP-based promoter analysis of selected HMGA2 target genes using the indicated antibodies and chromatin from MEF treated as in D. (F) DIP-based promoter analysis of selected HMGA2 target genes using antibodies specific for 5-methylcytosine (5mC) and genomic DNA from MEF treated as in D. (G) DIP-based promoter analysis of selected HMGA2 target genes using the indicated antibodies and genomic DNA from *Hmag2+/+* or *Hmag2-/-* MEF that were transfected with control (-) or *Gadd45a*-specific small interfering RNA (siRNA) as indicated. (H) ChIP-based promoter analysis of selected HMGA2 target genes using pH2A.X-specific antibodies and chromatin from MEF treated as in G. In all bar plots, data are shown as means ± s.e.m. (*n* = 3 independent experiments); asterisks, *P* values after *t*-Test, *** *P* ≤ 0.001; ** *P* ≤ 0.01; * *P* ≤ 0.05; ns, non-significant. See also Figure S6. The statistical test values of each plot are shown in Supplementary Table S3.

### Inhibition of the FACT complex counteracts fibrosis hallmarks in IPF

The clinical potential of the here proposed mechanism of transcription regulation (Figure 5B) was approached by placing it in the context of the most common interstitial lung disease, IPF, in which TGFB signaling plays a key role (Rubio et al., 2019). RNA-seq in primary human lung fibroblasts (hLF) isolated from control (*n* = 3) and IPF (*n* = 3) patients revealed high expression levels of *HMGA2*, *SUPT16H* and *SSRP1* in IPF patients when compared to control donors (Figure S7A). Further, cross-analysis of RNA-seq in primary hLF isolated from control and IPF patients (Rubio et al., 2019) with RNA-seq in *Hmga2+/+* MEF that were non-treated or treated with TGFB (Figure 7A) allowed us to identify 923 orthologue genes that were at least 1.5 fold significantly increased in IPF hLF and in TGFB treated MEF when compared to the corresponding control cells. Gene set enrichment analysis (GSEA) (Subramanian et al., 2005) based on normalized enrichment scores (NSE) revealed significant enrichment of orthologue genes related to EMT, TGFB1 signaling pathway, inflammatory response, MYC target genes, UV response, fatty acid metabolism, among others (Figure 7B). In addition, graphical representation of the enrichment profile showed high enrichment scores (ES) for EMT and TGFB1 signaling pathway as the top two items of the ranked list (Figure 7C). To determine the role of the HMGA2-FACT-ATM-pH2A.X axis in IPF we analyzed *GATA6*, *MTOR* and *IGF1* in Ctrl and IPF hLF (Figures 7D-F). Correlating with the results obtained in MEF after TGFB1 treatment (Figures 6D-F), we detected in IPF hLF increased expression of *GATA6*, *MTOR* and *IGF1* (Figure 7D), as well as increased levels of pH2A.X and H2A.X in their promoters (Figures 7E and S7B), whereas 5mC levels were reduced (Figure 7F). Strikingly, FACTin treatment counteracted the effects observed in IPF hLF, supporting the involvement of the HMGA2-FACT-ATM-pH2A.X axis in this interstitial lung disease. To test this hypothesis, we monitored various hallmarks of fibrosis in Ctrl and IPF hLF, such as expression of fibrotic markers by qRT-PCR (Figure 7G), levels of ECM proteins by Hydroxyproline and Sircol assays (Figures 7H, top and S7C), cell proliferation by bromodeoxyuridine (BrdU) incorporation assay (Figure 7H, middle) and cell migration by Transwell invasion assay followed by hematoxylin and eosin (H&E) staining (Figure 7H, bottom). Remarkably, FACTin treatment of IPF hLF significantly reduced all hallmarks of fibrosis analyzed, thereby suggesting the use of FACTin for therapeutic approaches against IPF. Moreover, our *in vitro* findings in primary hLF were also confirmed *ex vivo* using human precision-cut lung slices (hPCLS) from 3 different IPF patients (Figures 7I and S7D-G). FACTin treatment of IPF hPCLS reduced the levels of the fibrotic markers COL1A1, FN, smooth muscle actin alpha 2 (ACTA2), the mesenchymal marker vimentin (VIM), as well as HMGA2 and pH2A.X. In contrast, the levels of DNA-RNA hybrids were increased after FACTin treatment.

**Figure 7:**
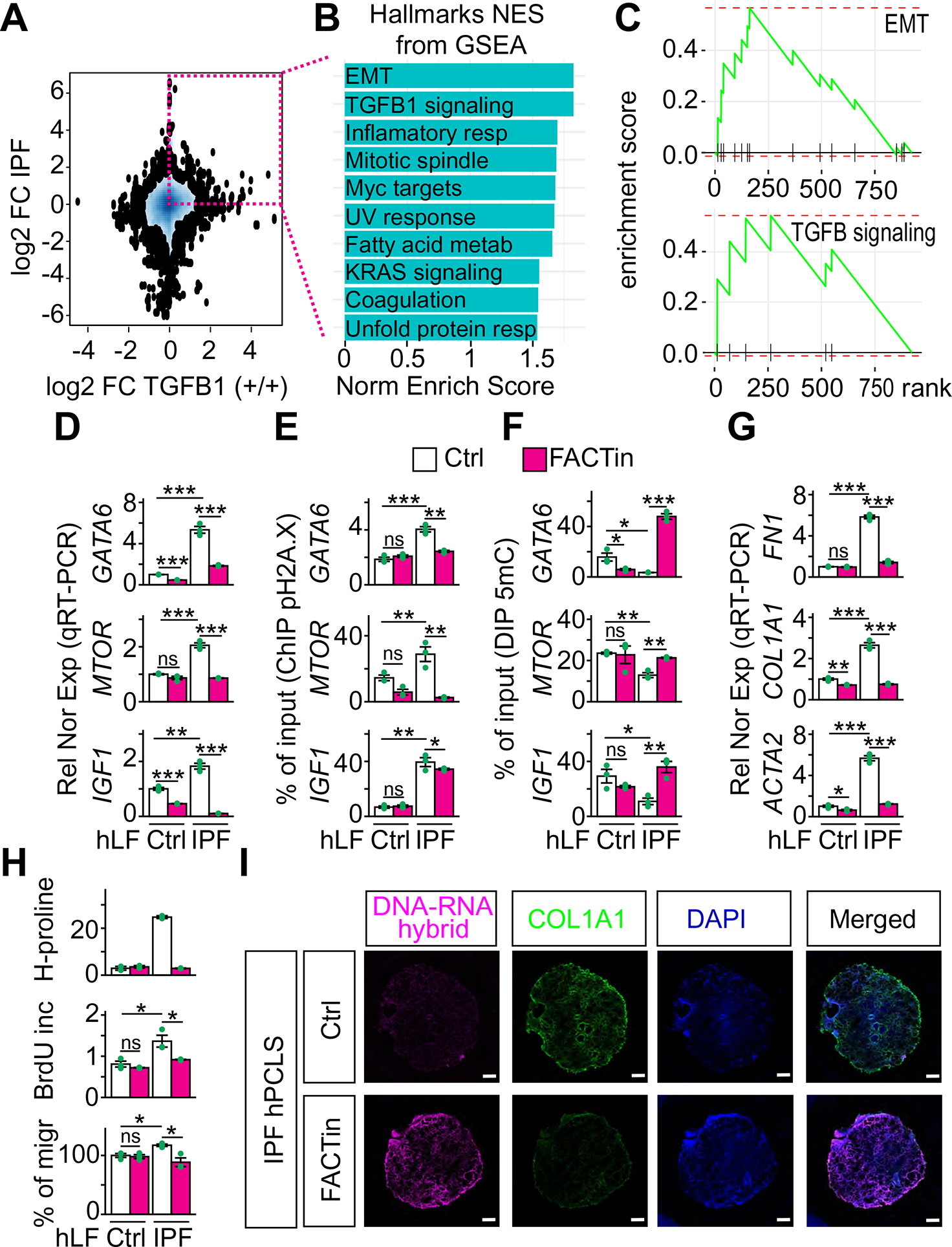
Inhibition of the FACT complex counteracts fibrosis hallmarks in IPF. (A) RNA-seq-based comparison of gene expression in IPF and after TGFB1 treatment. 2D Kernel Density plot representing the log2 fold change between gene expression in primary human lung fibroblasts (hLF) from IPF patients vs. control donors on the y-axis and log2 fold change between gene expression in *Hmga2+/+* MEF treated with TGFB1 vs. non-treated on the x-axis. Square, genes with log2 FC > 0.58 and *P* ≤ 0.05 in both, hLF IPF and TGFB1-treated MEF. *P* values after Wald test. (B) Gene set enrichment analysis (GSEA) using the normalized enrichement scores (NES) of genes inside the square in A. EMT, epithelial-mesenchymal transition; resp, response. (C) GSEA line profile of the top two enriched pathways in B. (D) qRT-PCR-based expression analysis of selected HMGA2 target genes in hLF from control donors (Ctrl) or IPF patients that were non-treated (Ctrl) or treated FACT inhibitor (FACTin; CBLC000 trifluoroacetate) as indicated. (E) ChIP-based promoter analysis of selected HMGA2 target genes using pH2A.X-specific antibodies and chromatin from hLF treated as in D. (F) DIP-based promoter analysis of selected HMGA2 target genes using 5mC-specific antibodies and genomic DNA from hLF treated as in D. (G) qRT-PCR-based expression analysis of fibrotic markers in hLF treated as in D. FN1, fibronectin; COL1A1, collagen; ACTA2, smooth muscle actin alpha 2. (H) Functional assays for IPF hallmarks in Ctrl or IPF hLF treated as in D. Top, hydroxyproline assay for collagen content. Middle, proliferation assay by BrdU incorporation. Bottom, Transwell invasion assay. (I) Representative pictures from confocal microscopy after immunostaining using the antibody S9.6 or COL1A1-specific antibody in human precision-cut lung slices (hPCLS) from IPF patients (*n* = 3 biologically independent experiments). The hPCLS were treated as in D. DAPI, nucleus. Scale bars, 500 μm. In all bar plots, data are shown as means ± s.e.m. (*n* = 3 independent experiments); asterisks, *P* values after *t*-Test, *** *P* ≤ 0.001; ** *P* ≤ 0.01; * *P* ≤ 0.05; ns, non-significant. See also Figure S7. The statistical test values of each plot are shown in Supplementary Table S3.

## DISCUSSION

Here we uncovered an unprecedented mechanism of transcription initiation of TGFB1-responsive genes mediated by the HMGA2-FACT-ATM-pH2A.X axis. The lyase activity of HMGA2 induces DNA nicks at the TSS, which are required by the FACT complex to incorporate nucleosomes containing H2A.X at specific positions relative to the TSS. The position of the first nucleosome containing H2A.X determines not only the basal transcription activity of the corresponding genes, but also the strength of their inducibility after TGFB1 treatment. Further, ATM-mediated phosphorylation of H2A.X at S139 is required for repair-mediated DNA demethylation and transcriptional activation. Our data support a sequential order of events, in which specific positioning of nucleosomes containing the classical DNA damage marker pH2A.X precedes DNA demethylation and transcription initiation, thereby supporting the hypothesis that chromatin opening involves intermediates with DNA breaks that require mechanisms of DNA repair that ensure the integrity of the genome.

The presented experimental data robustly support the molecular mechanism proposed here. Nevertheless, this work should be considered as starting point of future projects that will elucidate exciting questions that remained open. For example, it is currently unclear how HMGA2 is targeted to specific promoters. One answer to this question might be the inducibility of specific genes in determined signaling pathways, including TGFB, IGF and WNT signaling pathways, especially since they have been related to HMGA2 (Brants et al., 2004; Li et al., 2012; Singh et al., 2014; Singh et al., 2015). However, this is just a partial answer, since there are genes that are inducible by these signaling pathways in an HMGA2-independent manner, as the ones shown in clusters 2 and 4 of Figure 5A, as well as there are HMGA2 target genes that are not induced by these signaling pathways. Interestingly, we have observed that 46% of the genes inside the top 15% candidates show nascent ncRNAs in antisense orientation (Figure S4H, top). In addition, 38% of the genes inside the top 15% candidates contain GC skews that favor the formation of R-loops (Figure S4H, bottom). The DNA-ncRNA hybrids of the R-loops might be an excellent option to tether HMGA2 to specific promoters. Similarly, triple-helical RNA-DNA-DNA structures (triplex) at enhancers and promoters allow lncRNAs to recruit protein complexes to specific genomic regions and regulate gene expression (Blank-Giwojna et al., 2019; Kuo et al., 2019). Interestingly, Summer and colleagues demonstrated in a seminal work that HMGA2 binds and efficiently cleaves ssDNA containing abasic sites *in vitro* (Summer et al., 2009). It will be the scope of our future work to determine whether the binding affinity and the cleavage efficiency of HMGA2 increase when the DNA substrate for the intrinsic lyase activity of HMGA2 is part of a DNA-RNA hybrid in an R-loop. Supporting this line of ideas, Arab and colleagues recently reported in a pioneering work that GADD45A preferentially binds DNA-RNA hybrids and R-loops rather than ssDNA, dsDNA or RNA (Arab et al., 2019). Strikingly, we have shown that loss of HMGA2 lyase activity increased R-loop levels (Figures 4F-G), while binding of GADD45A to R-loops was reduced (Figures 5G-H), thereby supporting the hypothesis that single-strand breaks of the DNA moiety in the DNA-RNA hybrids are required for GADD45A binding. Another open question, from the mechanistic point of view, is related to the origin of the abasic sites that are bound and cleaved by HMGA2. Since 38% of the genes inside the top 15% candidates contain GC skews and active DNA demethylation is part of the mechanism of transcription initiation proposed here, a plausible explanation for the generation of abasic sites is the participation of DNA glycosylases that removes the bases from 5-methycytosine (5mC) or thymine as deamination product of 5mC. In addition, HMGA2 has been shown to interact with APEX1 (apuryinic/apyrimidinic site endonuclease 1) (Summer et al., 2009) and XRCC6 (X-ray repair cross-complementing protein 6, a ssDNA-dependent helicase also known as Ku70) (Sgarra et al., 2008), thereby supporting the involvement of the base excision repair (BER) machinery during HMGA2 function. It will be interesting to perform functional experiments demonstrating the requirement of the BER machinery during the mechanism of transcription initiation proposed here. Another interesting aspect will be to elucidate the molecular mechanism regulating transcription of the genes inside of the TGFB cluster 3 (Figure 6A), which are TGFB inducible in *Hmga2+/+* MEF but they become TGFB repressible after *Hmga2*-KO.

We have recently reported a mechanism of transcription repression mediated by the multicomponent RNA–protein complex MiCEE (Singh et al., 2018). In addition, we have shown that in IPF reduced levels of the micro RNA lethal 7d (*MIRLET7D*, also known as *let-7*) and hyperactive EP300 compromise the epigenetic gene silencing mediated by the MiCEE complex (Rubio et al., 2019). The results presented here strongly imply an opposite function of the MiCEE complex and the HMGA2-FACT-ATM-pH2A.X axis in the context of TGFB1 signaling and IPF. Supporting this line of evidence, it has been reported that *MIRLET7D* targets *HMGA2* mRNA, thereby preventing TGFB1-induced EMT and renal fibrosis (Wang et al., 2016). Further, reduction of mature *MIRLET7* levels by the oncofetal protein LIN28B allows HMGA2 to drive an epigenetic program during pancreatic ductal adenocarcinoma (PDAC), one of the most lethal malignancies. (Kugel et al., 2016). It will the scope of our future work to confirm the opposite function of the MiCEE complex and the HMGA2-FACT-ATM.pH2A.X axis within the context of fibrosis in different organs, including lung, kidney and liver.

We demonstrated the biological relevance of our data within the context of TGFB1 signaling (Figures 6 and S6). TGFB1 treatment induced promoter specific increase of pH2A.X and pPol II, whereas 5mC levels were decreased, resulting in transcription activation in *Hmga2*- and FACT-dependent manner (Figures 6A-F). Interestingly, *Gadd45a*-LOF interfered with the 5mC decrease (Figure 6G), without affecting pH2A.X levels (Figure 6H), supporting the sequential order of events proposed here (Figure 5B), in which GADD45A acts downstream of the HMGA2-FACT-ATM-pH2A.X axis. Consistent with our findings, Thillainadesan and colleagues reported TGFB induced active DNA demethylation and expression of the *p15^ink4b^* tumor suppressor gene (Thillainadesan et al., 2012). While published reports showed the effect of TGFB on specific genes (Duan and Derynck, 2019; Singh et al., 2015; Thillainadesan et al., 2012), in this report we demonstrated the genome wide effect of TGFB treatment affecting the global nuclear architecture and strongly suggesting future NGS studies. Following a similar line of ideas, Negreros and colleagues recently reported genome wide changes on DNA methylation induced by TGFB1 (Negreros et al., 2019). The translational potential of our work was demonstrated within the context of IPF (Figures 7 and S7), in which TGFB1 signaling plays an important role. Inhibition of the HMGA2-FACT-ATM-pH2A.X axis reduced all fibrotic hallmarks *in vitro* (using primary hLF) and *ex vivo* (using hPCLS). Interestingly, the FACT complex is a potential marker of aggressive cancers with low survival rates (Garcia et al., 2013) and FACTin is being tested in a clinical trial for cancer treatment (ClinicalTrials.gov Identifier: NCT01905228, NCT02931110). Our work provides the molecular basis for future studies developing therapies against IPF using FACTin.

## ACKNOWLEDGMENTS

We thank Roswitha Bender for technical support. Guillermo Barreto is funded by the „Université Paris Est” (UPEC, Créteil, France), the Max-Planck-Society (MPG, Munich, Germany) and the „Deutsche Forschungsgemeinschaft” (DFG, Bonn, Germany) (BA 4036/4-1). Indrabahadur Singh is funded by the DFG (Bonn, Germany) through Emmy Noether program (SI 2620/1-1). Rafael Castillo received a doctoral fellowship from CONACyT (2019-000003-01EXTF-00156, Mexico). Jürgen Bernhagen acknowledges support from DFG under Germany’s Excellence Strategy within the framework of the Munich Cluster for Systems Neurology (EXC 2145 SyNergy – ID 390857198) and within the LMU-EXC strategic cooperations program LMU/Singapore. This work was done according to the program of competitive growth of the Kazan Federal University and the Russian Government. Biomaterials were provided by the European IPF Registry and Biobank (eurIPFreg).

## AUTHOR CONTRIBUTIONS

SD, KR, IS, SG, JG, JC, RCN, PB, AM, MBH and GB designed and performed the experiments. HCF, JB, CMC, SB, AG, KTP, GD, MW, TB and DPG were involved in study design. GB, SD, KR, IS, SG, JG and JC designed the study, analyzed the data. GB, SD, KR and IS wrote the manuscript. All authors discussed the results and commented on the manuscript.

## STAR METHODS

### Key Resources Table

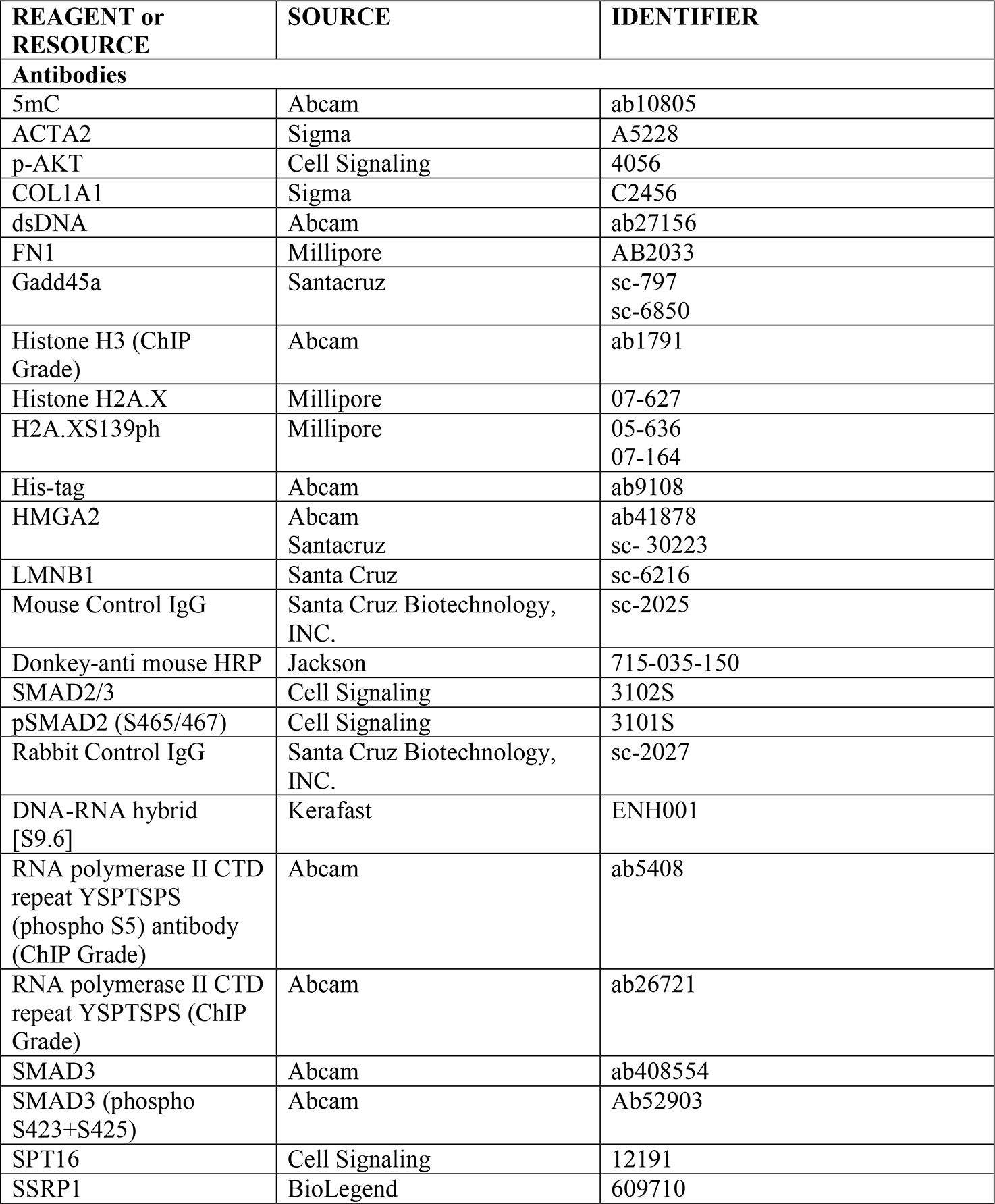
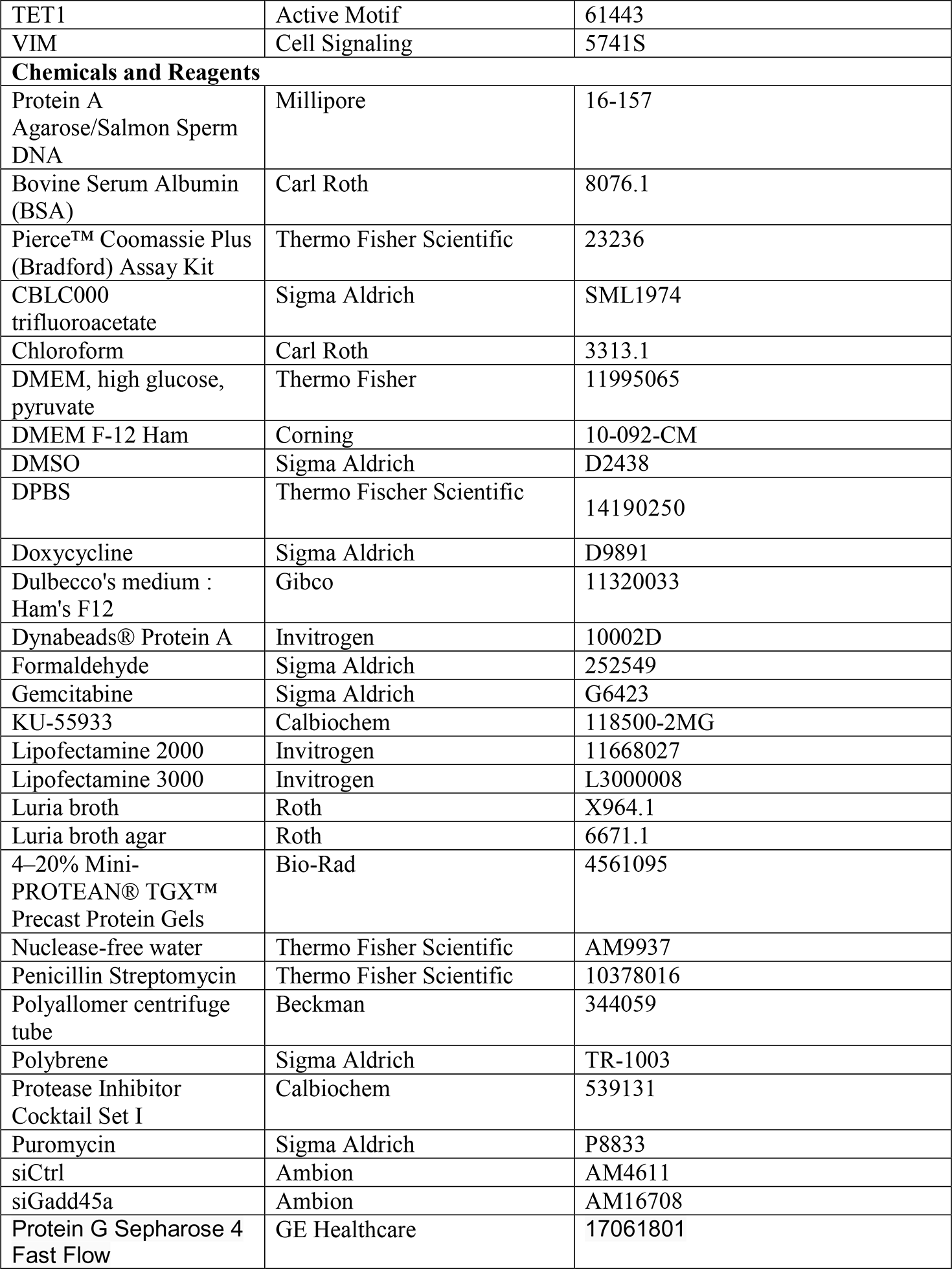
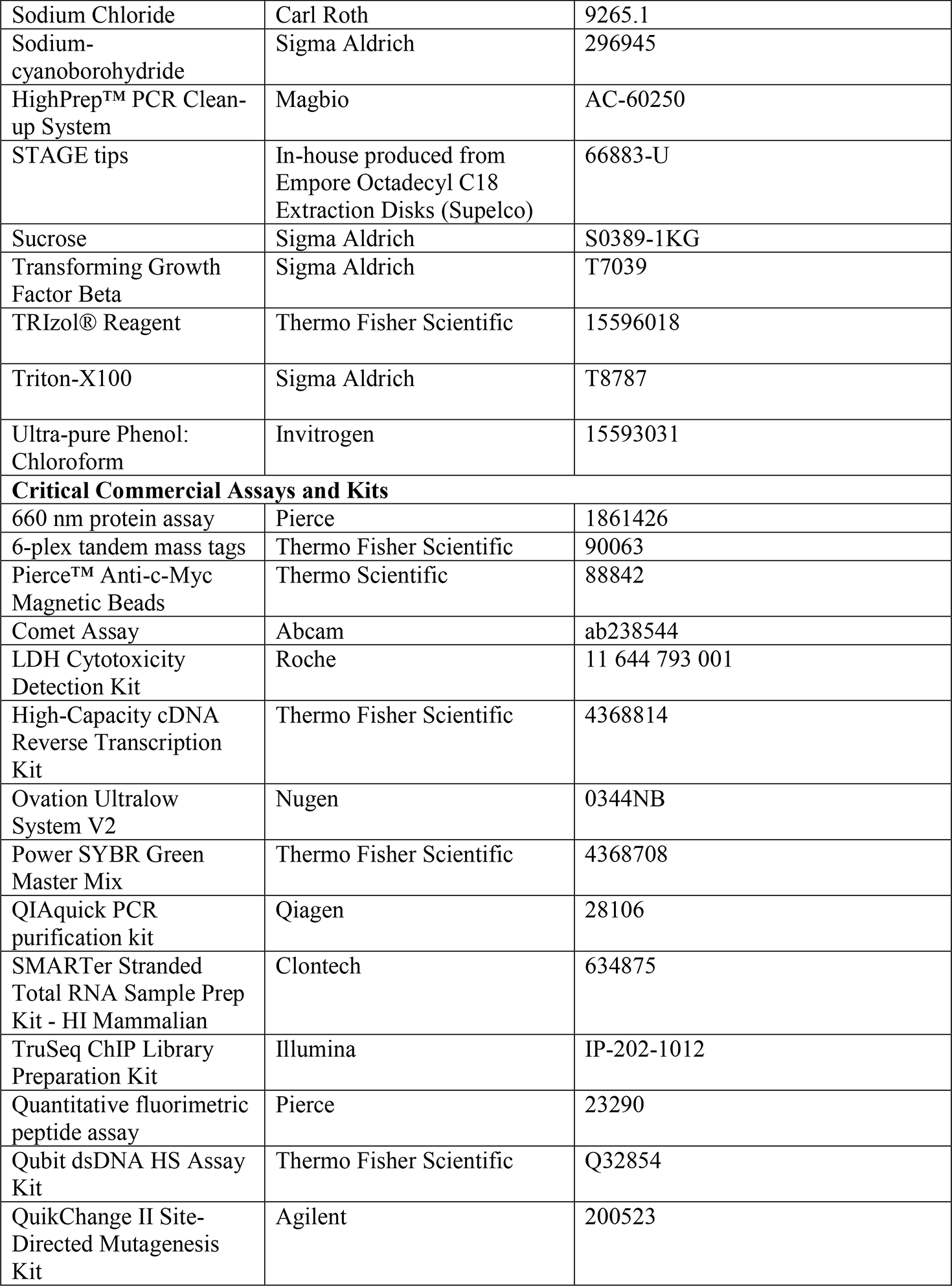
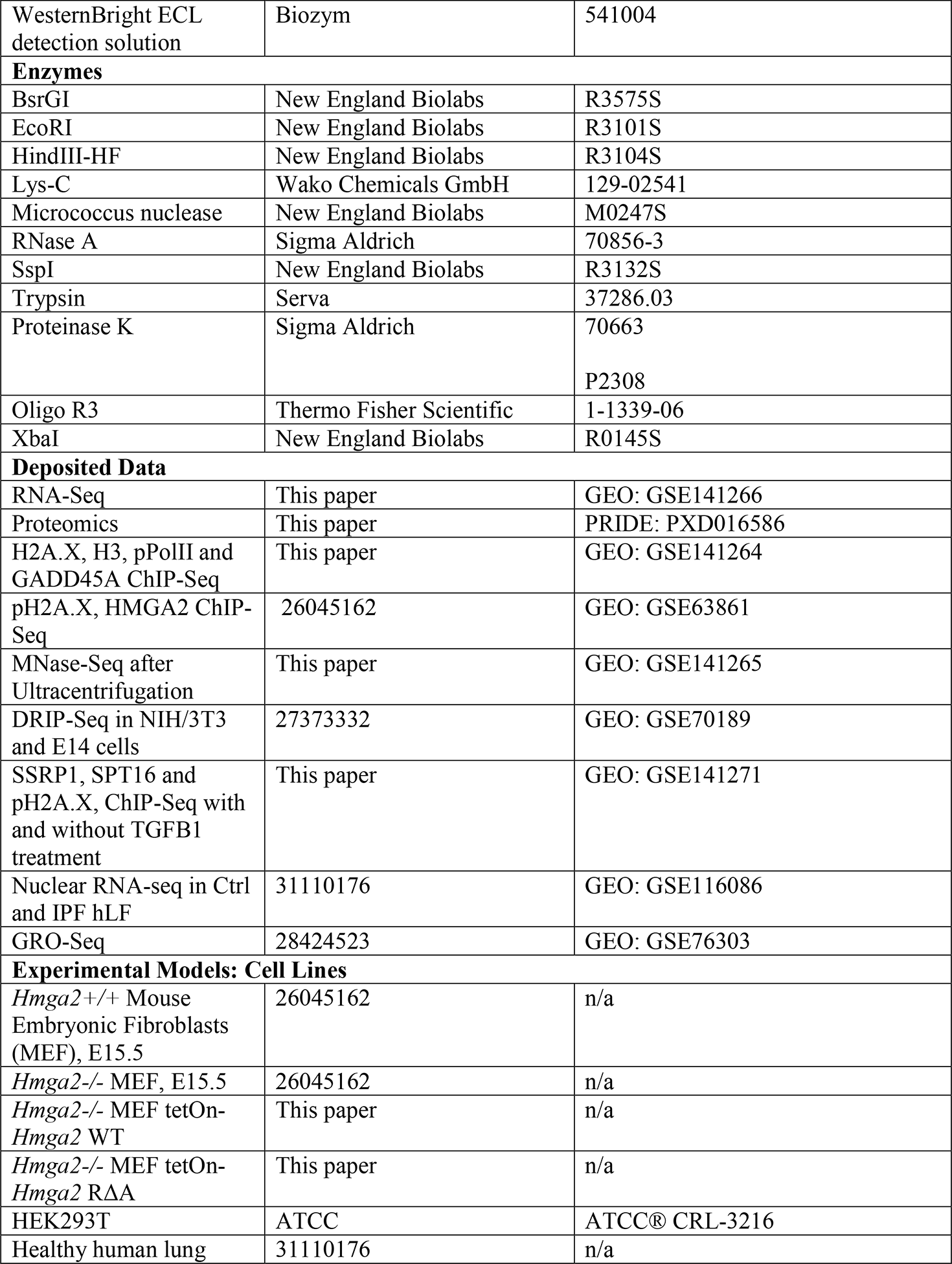
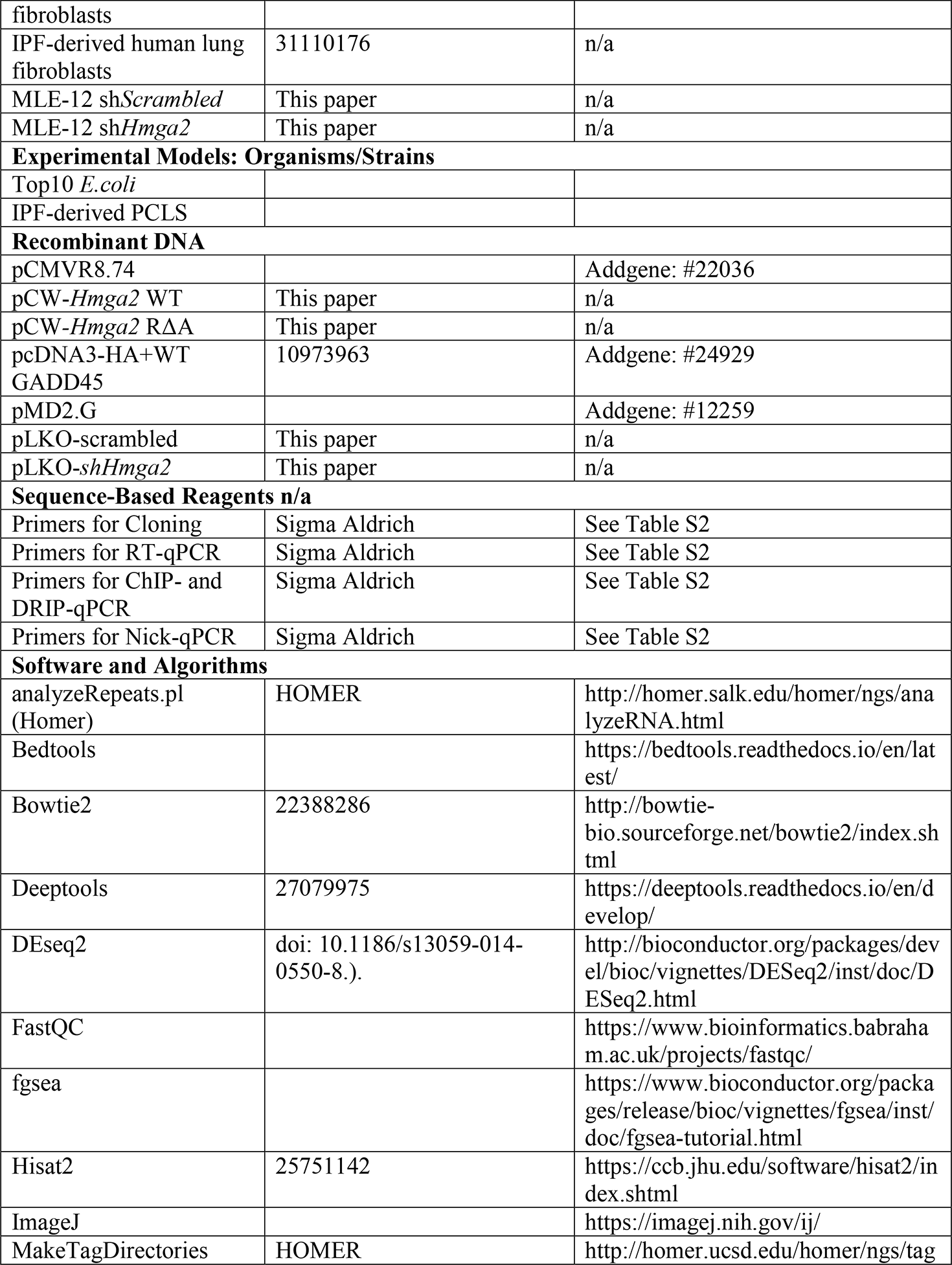
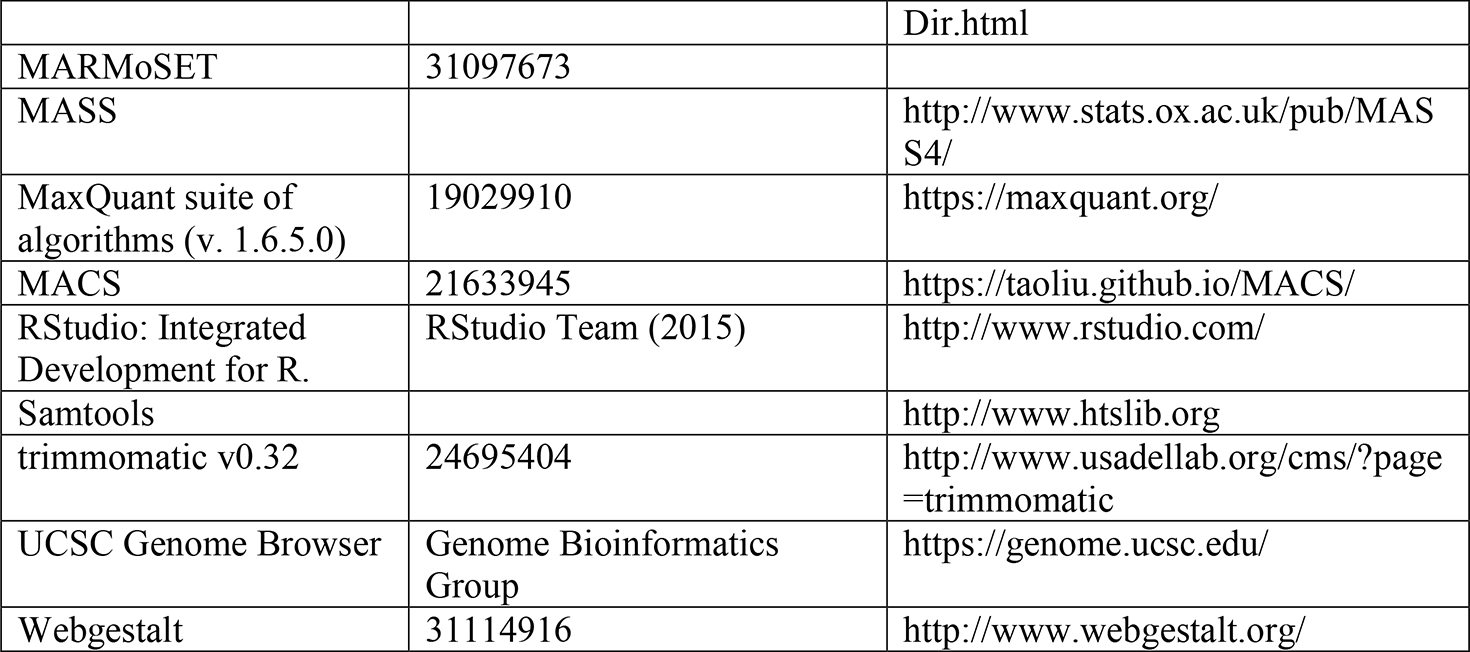

## Contact for Reagent and Resource Sharing

Please direct any requests for further information or reagents to Lead Contact Guillermo Barreto, Brain and Lung Epigenetics (BLUE), Laboratoire Croissance, Réparation et Régénération Tissulaires (CRRET), Université Paris Est Créteil (UPEC), F-94000, Créteil, France.

## Study design

This study was performed according to the principles set out in the WMA Declaration of Helsinki; the underlying protocols were approved by the ethics committee of Medicine Faculty of the Justus Liebig University in Giessen, Germany (AZ.111/08-eurIPFreg) and the Hannover Medical School (no. 2701-2015). In this line, all patient and control materials were obtained through the UGMLC Giessen Biobank (member of the DZL Platform Biobanking) and the Biobank from the Institute for Pathology of the Hannover Medical School as part of the BREATH Research Network. We used anonymized patient material.

## Experimental Model and Subject Details

### Cell culture

All studies were done on immortalized MEF cultivated for less than twenty passages. *Hmga2* wild type (+/+) and knockout (-/-) primary mouse embryonic fibroblast (MEF) were prepared and used as described earlier (Singh et al., 2015). These cells were immortalized using SV40. MEF and Human embryonic kidney cell HEK293T (ATCC, CRL-11268) were cultured at 37 °C in 5% CO_2_ in DMEM medium with 4.5 g/l glucose, 10% FCS 4 mM L-Glutamine, 1 mM Pyruvate, 100 U/ml penicillin and 100 U/ml streptomycin. Mouse lung epithelial cells (MLE-12, ATCC CRL-2110) were cultured in Dulbecco’s Modified Eagle Medium: Ham’s F-12 Nutrient Mixture (5% FCS, 100 U/ml penicillin and 100 U/ml streptomycin) at 37 °C in 5% CO_2_. Cells were 1x PBS washed, trypsinized with 0.25% (w/v) Tryspin and subcultivated at the ratio of 1:5 to 1:10. Primary fibroblast from Ctrl and IPF patients were cultured in complete MCDB131 medium (8% FCS, 1% L-glutamine, penicillin 100 U/ml, streptomycin 0.1 mg/ml, EGF 0.5 ng/ml, bFGF 2 ng/ml, and insulin 5 μg/ml)) at 37 °C in 5% CO_2_. Because of the concern that the phenotype of the cells is altered at higher passage, cells between passages 4 and 6 were utilized in the experiments described here. During subculturing, cells were washed with 1x PBS, trypsinized with 0.25% (w/v) trypsin and subcultivated at the ratio of 1:5 to 1:10.

### Cell treatments, transfections and siRNA-mediated knockdown

MEF were treated with 1 μg/ml doxycycline or DMSO (used as solvent for doxycycline) for 4, 6 or 24 h to induce the expression of transgenes. Initial FACT complex and ATM kinase inhibition was performed with 5 μM CBLC000 trifluoroacetate (FACTin, Sigma Aldrich) for 2 h or 1 μM KU-55933 (ATMi, Calbiochem) for 6 h, respectively. MEF were transiently transfected either with 100 nM siCtrl (negative control; AM4611, Ambion) or *siGadd45a* (siG45; AM16708, Ambion) for 48 h. TGFB1 signaling was induced with 10 ng/ml human recombinant TGFβ1 (Sigma Aldrich) and chromatin changes were assayed after 3 h and gene expression alterations after 24 h incubations.

Primary fibroblast from Ctrl and IPF patients were cultured in complete MCDB131 medium (8% FCS, 1% L-glutamine, penicillin 100 U/ml, streptomycin 0.1 mg/ml, EGF 0.5 ng/ml, bFGF 2 ng/ml, and insulin 5 μg/ml) at 37 °C in 5% CO_2_. Because of the concern that the phenotype of the cells is altered at higher passage, cells between passages 4 and 6 were utilized in the experiments described here, except cells for screening were used at earlier passages. During subculturing, cells were washed with 1X PBS, trypsinized with 0.25% (w/v) trypsin and subcultivated at the ratio of 1:5 to 1:10. For IPF resolution experiments, primary hLF were treated with 5 μM FACTin for 12 h.

### Bacterial culture

For cloning experiments, chemically competent *E. coli* TOP10 (Thermo Fisher Scientific) were used for plasmid transformation. TOP10 strains were grown in Luria broth (LB) at 37 °C with shaking at 180 rpm on LB agar at 37 °C overnight.

## Method Details

### Chromatin Immunoprecipitation (ChIP)

Cells were cross-linked by 1% formaldehyde for 10 min, lysed, and sonicated with Diagenode Bioruptor to an average DNA length of 300-600 bp. After centrifugation, the soluble chromatin was immunoprecipitated using antibodies specific for H3 (Abcam), H4 (Abcam), H2A (Abcam), H2B (Abcam), pH2A.X (Millipore), H2A.X (Millipore), HMGA2 (Abcam), pPol II (Abcam), GADD45A (Santacruz), TET1 (Active Motif) and IgG as a control (Santa Cruz Biotechnology). Precipitated chromatin complexes were removed from the beads by incubating with 50 μl of 1% SDS, 0.1 M NaHCO3, for 30 min while vortexing every 5 min. Reverse cross-linked immunoprecipitated chromatin was purified using the QIAquick PCR purification kit (Qiagen) and subjected to qPCR or next-generation sequencing. For qPCR, the percentage of input was calculated after subtracting the IgG background, if not stated elsewhere.

### ChIP sequencing and data analysis in *Hmga2+/+* and *Hmga2-/-* MEF

Libraries were prepared according to Illumina’s instructions accompanying the Ovation Ultra Low Kit. Single-end sequencing was performed on an Illumina HiSeq2500 machine at the Max Planck-Genome-Centre Cologne. Raw reads were visualized by FastQC to determine the quality of the sequencing. Trimming was performed using trimmomatic v0.32 with the following parameters LEADING:3 TRAILING:3 SLIDINGWINDOW:4:15 MINLEN:50 CROP:63 HEADCROP:13. High quality reads were mapped by using Bowtie2 to mouse genome mm10. ChIP-seq data were represented as aggregate plot or heat maps using deeptools following their instructions. Bam-files were converted to Bed-files using Bedtools’ bamToBed command. Genome browser snapshots were created with Homer using makeTagDirectory and makeUCSCfile. Reads were normalized to 30 million reads. Peak calling was performed by using model-based analysis for ChIP-seq (MACS) (Feng et al., 2011) with a cut-off of p<0.01 and the following parameters: --nonmodel, –shift size 30, and an effective genome size -g of 1.87e9. Peaks were annotated by using annotePeaks.pl for mm10 from Homer. From the 200,051 peaks found for pH2A.X in *Hmga2+/+* MEF, 3,935 peaks were annotated in the promoter-TSS region and were used for further analysis.

### Identification of position clusters and analysis of inducibility

The enrichment of pH2A.X at TSS plus 250 bp downstream in the top 15% candidates was clustered using the *k*-means algorithm implemented in deeptools’ plotHeatmap command. From the 3 position clusters identified, genes in *Hmga2+/+* MEF with a FC more than 1.5 to *Hmga2-/-* were selected. For expression analysis, RPKMs less than 10 were included.

### RNA isolation, reverse transcription, quantitative PCR

Total RNA was isolated with Trizol (Invitrogen) and quantified using a Nanodrop Spectrophotometer (ThermoFisher Scientific, Germany). Synthesis of cDNA was performed using 0.5-1 µg total RNA and the High Capacity cDNA Reverse Transcription kit (Applied Biosystems). Quantitative real-time PCR reactions were performed using SYBR® Green on the Step One plus Real-time PCR system (Applied Biosystems). The housekeeping genes *Tuba1a* and *HPRT1* was used to normalize gene expression (Mehta et al., 2015). Primer pairs used for gene expression analysis are described in Supplemental Information, Table S2.

### Native chromatin fractionation and sucrose gradient ultracentrifugation

Two 15 cm dishes with MEF were washed with 1x PBS and pellets were resuspended in 1 ml of lysis buffer (10 mM HEPES pH 7.4, 10 mM KCl, 0.05% NP-40, 1 mM DTT, 25 mM NaF, 0.5 mM Na_3_VO_4_, 40 μg/ml phenylmethylsulfonyl fluoride and protease inhibitor). After incubating 20 min on ice, cells were spun down at 300 x g at 4 °C for 10 min. The nuclei were washed once in lysis buffer and were then resuspended in 2 volumes of Low Salt Buffer (10 mM Tris-HCl pH 7.4, 0.2 mM MgCl_2_ supplemented with protease and phosphatase inhibitors) including 1% Triton-X100. After 15 min incubation on ice, cells were spun down at 300 x g at 4 °C for 10 min. The pellet was washed in 1 ml MNase digestion buffer (10 mM Tris-HCl, 25 mM NaCl, 1 mM CaCl2, 1 mM DTT, 25 mM NaF, 0.5 mM Na_3_VO_4_, 40 μg/ml phenylmethylsulfonyl fluoride and protease inhibitor) and resuspended again in 1 ml MNase digestion buffer with 1,250 Units MNase (NEB Biolabs). Chromatin-MNase mix was incubated at 37 °C for 30 min. MNase reaction was stopped by adding 50 mM EDTA. Samples were sonicated for 30 sec on / 30 sec off using the Bioruptor with high amplitude. Chromatin was spun down at 14,000 rpm at 4 °C for 10 min and the supernatant was used for ultracentrifugation. Sucrose gradients (5% to 40%) were prepared in 1,800 μl low salt buffer (10 mM NaCl, 10 mM Tris-HCl pH7.4, 0.2 mM EDTA, 0.2 mM DTT, 20 mM NaF, 20 mM Na_3_VO_4_, 40 μg/ml phenylmethylsulfonyl fluoride and protease inhibitor) in polyallomer centrifuge tube (Beckman). Fragmented native chromatin was loaded on top of the 9 ml 5% to 40% sucrose gradients and centrifuged for 16 h and 30 min at 37,500 rpm in a SW50.1 ultracentrifuge rotor (Beckman Coulter). Following centrifugation, 11 fractions (1000 μl each) were collected manually from the bottom of the tubes. Later, these fractions were used for western blot, mass spectrometry and NGS.

Western blotting was performed using standard methods and antibodies specific for HMGA2 (Abcam), pPol II (Abcam), total Pol II (Abcam), SUPT16 (Cell Signaling), SSRP1 (Biolegend), pH2A.X (Millipore), H2A.X (Abcam), H2A (Abcam), H2B (Abcam) and H3 (Abcam) were used. Immunoreactive proteins were visualized with the corresponding HRP-conjugated secondary antibodies (Jackson) using the WesternBright ECL detection solutions (Biozym). Signals were detected and analyzed with Luminescent Image Analyzer (Las 4000, Fujifilm.

### Mass spectrometry: sample preparation, methods and data analysis

Proteins were methanol/chloroform precipitated from sucrose gradient fractions and dried pellets reconstituted in 8 M urea (Wessel and Flugge, 1984). Per fraction, 140 µg of protein (according to the 660 nm protein assay, Pierce), were subjected to in-solution digest using protein to enzyme ratios of 1:100 and 1:50 for Lys-C (Wako Chemicals GmbH) and trypsin (Serva), respectively (Graumann et al., 2008). The resulting peptide mixture was desalted and concentrated using Oligo R3 (Thermo Fisher Scientific) extraction (Billing et al., 2015). Peptides (5 µg according to the quantitative fluorimetric peptide assay, Pierce), were subsequently labeled using 6-plex tandem mass tags (Thermo Fisher Scientific) following the manufacturer’s protocol but employing a reagent to peptide ratio of four. Labeling channels were used for fractions from a replicate sucrose gradient as well as an internal standard sample consisting from an analogously treated mix of all replicate gradient input samples. After validation of labeling efficiency by liquid chromatography-tandem mass spectrometry (LC-MS2), samples were mixed by equal protein amount and 4 µg total peptides purified as well as concentrated using STAGE tips (Rappsilber et al., 2003). The subsequent LC-MS2 analysis of 50% of that peptide material used an in-house packed 70 μm ID, 15 cm reverse phase column emitter (ReproSil-Pur 120 C18-AQ, 1.9 μm, Dr. Maisch GmbH) with a buffer system comprising solvent A (5% acetonitrile, 0.1% formic acid) and solvent B (80% acetonitrile, 0.1% formic acid). Relevant instrumentation parameters are extracted using MARMoSET and included in the supplementary material (Kiweler et al., 2019). Peptide/protein group identification and quantitation was performed using the MaxQuant suite of algorithms (v. 1.6.5.0) against the mouse uniprot database (canonical and isoforms; downloaded on 2019/01/23; 86695 entries) (Cox and Mann, 2008; Cox et al., 2011). For downstream analysis, intensities of fractions were divided by their corresponding inputs and samples that were divided by zero were set to 0.1.

### MNase-sequencing and data analysis

For DNA purification from sucrose ultracentrifugation fractions, 200 µl of fractions were resuspended with 200 µl 1x PBS and incubated with 0.5 µl RNase A (10 mg/ml, Sigma Aldrich) for 15 min at 37 °C. Samples were resuspended with 400 µl Ultra-pure phenol:chloroform (Invitrogen) and incubated for 5 min at RT. After centrifugation for 5 min at 14,000 rpm at 4 °C, the clear phase containing DNA was transferred into a fresh tube. 40 µl of 3 M sodium acetate pH 4.9 and 1 ml of ethanol were added to the samples and DNA was precipitated for 30 min at −80 °C. DNA-mix was spun down for 30 min at 14,000 rpm at 4 °C and the DNA pellets were washed with 70% ice cold ethanol. After centrifugation for 15 min at 14,000 rpm at 4 °C, the pellets were dried at RT and further resuspended in 50 µl of nuclease-free water and heated for 15 min at 37 °C. DNA was analyzed by agarose gel electrophoresis or NGS.

For sequencing, purified DNA was quantified by Qubit dsDNA HS Assay Kit (Thermo Fisher Scientific). 10 ng DNA was used as input for TruSeq ChIP Library Preparation Kit (Illumina) with following modifications. Instead of gel-based size selection before final PCR step, libraries were size selected by SPRI-bead based approach after final PCR with 18 cycles. In detail, samples were first cleaned up by 1x bead:DNA ratio to eliminate residuals from PCR reaction, followed by 2-sided-bead cleanup step with initially 0.6x bead:DNA ratio to exclude larger fragments. Supernatant was transferred to new tube and incubated with additional beads in 0.2x bead:DNA ratio for eliminating smaller fragments, like adapter and primer dimers. Bound DNA samples were washed with 80% ethanol, dried and resuspended in TE buffer. Library integrity was verified with LabChip Gx Touch 24 (Perkin Elmer). Sequencing was performed on the NextSeq500 instrument (Illumina) using v2 chemistry with 2×38bp paired setup. Raw reads were visualized by FastQC to determine the quality of the sequencing. Trimming was performed using trimmomatic with the following parameters LEADING:3 TRAILING:3 SLIDINGWINDOW:4:15 HEADCROP:5, MINLEN:15. High quality reads were mapped by using with bowtie2 to mouse genome mm10. For downstream analysis, fragments between 100 and 200 bp were selected and the reads were centered. Reads were normalized by an RPKM measure and represented as log2 enrichment over the corresponding inputs. To avoid division through zero, zero counts were pseudo-counted as “1”.

### Generation of HMGA2 lyase mutant, doxycycline inducible MEF and stable knockdown of

#### *Hmga2* in MLE-12 cells

The lyase deficient mutant of HMGA2 was generated by sequential site-directed mutagenesis (QuikChange II Site-Directed Mutagenesis Kit, Agilent) of the first three arginines in the third AT-Hook domain. Mutagenesis primers for *Hmga2* are listed in Table S2. All constructs were sequence verified.

Replication-deficient lentiviruses containing doxycycline inducible Ctrl (Empty), *Hmga2* WT-myc-his, *Hmga2* RΔA-myc-his, pLKO-*scrambled* or pLKO-*shHmga2* were produced by transient transfection of pCMVR8.74 (Addgene: #22036), pMD2.G (Addgene: #12259) and transfer plasmid into HEK293T cells in a 6-well plate. Viral supernatants were collected after 48 h, spun down at 4,000 rpm for 20 min, and then used to transduce immortalized MEF in the presence of polybrene (10 μg/ml, Sigma). Forty-eight h later, MEF and MLE-12 cells were selected by stepwise increase with 1.5 to 3.0 or 4.0 μg/ml puromycin respectively and the pooled populations were used for various experiments.

#### Co-immunoprecipitation (Co-IP) and Western blot

MEF were washed three times in cold 1x PBS and scraped in 10 ml of lysis buffer (10 mM HEPES pH 7.4, 10 mM KCl, 0.05% NP-40, 1 mM DTT, 25 mM NaF, 0.5 mM Na_3_VO_4_, 40 μg/ml phenylmethylsulfonyl fluoride and protease inhibitor). Cells were incubated on ice for 20 min and then spun down at 300 x g for 10 min at 4 °C. Nuclear cell pellets were resuspended in 300 μl Co-IP buffer (50 mM Tris-HCl pH 7.4, 170 mM NaCl, 20% glycerol, 15 mM EDTA, 0.1% (v/v) Triton X-100, 20 mM NaF, 20 mM Na_3_VO_4_, 40 μg/ml phenylmethylsulfonyl fluoride and protease inhibitor) and were sonicated 5 times using the Bioruptor (30 sec on, 30 sec off). Soluble chromatin and proteins were collected after centrifugation at 10,000 x g at 4 °C for 10 min. Precleared nuclear protein lysates (500 μg) were incubated with 20 μl of anti-c-MYC tag antibody coupled to magnetic beads (Thermo Scientific). After two hours, beads were collected and washed 4 times with 500 μl ice-cold washing buffer (50 mM Tris-HCl pH 7.4, 170 mM NaCl, 15 mM EDTA, 0.4% (v/v) Triton X-100, 20 mM imidazole, 20 mM NaF, 20 mM Na_3_VO_4_, 40 μg/ml phenylmethylsulfonyl fluoride and protease inhibitor). Proteins were eluted in 30 µl 2 x SDS sample loading buffer while boiling at 95 °C for 5 min. Beads were removed and protein eluates were loaded on SDS-PAGE for western blot analysis. Western blotting was performed using standard methods and antibodies specific for SUPT16 (Cell Signaling), SSRP1 (Biolegend), H2A.X (Millipore) and HIS-tag (Abcam) were used. Immunoreactive proteins were visualized with the corresponding HRP-conjugated secondary antibodies (Jackson) using the WesternBright ECL detection solutions (Biozym). Signals were detected and analyzed with Luminescent Image Analyzer (Las 4000, Fujifilm). Protein concentrations were determined using Bradford kit (Pierce).

#### Preparation of nuclear, chromatin and nucleoplasm extracts

Cells were washed with 1x PBS and pellets were resuspended in 2 volumes of lysis buffer (10 mM HEPES pH 7.4, 10 mM KCl, 0.05% NP-40, 1 mM DTT, 25 mM NaF, 0.5 mM Na_3_VO_4_, 40 μg/ml phenylmethylsulfonyl fluoride and protease inhibitor (Calbiochem)). After 20 min incubation on ice, cells were spun down at 300 x g at 4 °C for 10 min. The supernatant was removed and nuclei were resuspended in whole cell lysate buffer (1% SDS, 10 mM EDTA pH 8, 50 mM Tris-HCl pH 8, 0.05% NP-40, 1 mM DTT, 25 mM NaF, 0.5 mM Na_3_VO_4_, 40 μg/ml phenylmethylsulfonyl fluoride and protease inhibitor). To obtain soluble nuclear proteins, resuspended nuclei were sonicated 5 times 30 sec on followed by 30 sec off using the Bioruptor (Diagenode). Insoluble proteins were removed by centrifugation for 10 min at 14,000 rpm at 4 °C.

For chromatin and nucleoplasm preparations, the protocol was adapted with minor modifications from (Sebastian et al., 2012). Cells were washed with 1 x PBS and pellets were resuspended in 2 volumes of lysis buffer (10 mM HEPES pH 7.4, 10 mM KCl, 0.05% NP-40, 1 mM DTT, 25 mM NaF, 0.5 mM Na_3_VO_4_, 40 μg/ml phenylmethylsulfonyl fluoride and protease inhibitor). After 20 min incubation on ice, cells were spun down at 300 x g at 4 °C for 10 min. The supernatant contains the cytoplasmic proteins. The nuclei were washed once in lysis buffer and were then resuspended in 2 volumes of Low Salt Buffer (10mM Tris-HCl pH7.4, 0.2 mM MgCl_2_ supplemented with protease and phosphatase inhibitors including 1% Triton-X100). After 15 min incubation on ice, cells were spun down at 300 x g at 4 °C for 10 min. The supernatant contains the nucleoplasm proteins and the pellet contains the chromatin. Pellets were resuspended in 2 volumes of 0.2 N HCl and incubated on ice for 20 min. Resuspended pellets were spun down at 14,000 rpm at 4 °C for 10 min and the supernatant containing acid soluble proteins was neutralized with the same volume of 1 M Tris-HCl pH 8.0. Western blotting was performed using standard methods and antibodies specific for SUPT16 (Cell Signaling), SSRP1 (Biolegend), HMGA2 (Abcam), H3 (Abcam) and AKT (Cell signaling) were used. Immunoreactive proteins were visualized with the corresponding HRP-conjugated secondary antibodies (Jackson) using the WesternBright ECL detection solutions (Biozym). Signals were detected and analyzed with Luminescent Image Analyzer (Las 4000, Fujifilm). Protein concentrations were determined using Bradford kit (Pierce).

### DNA-RNA hybrid immunoprecipitation (DRIP)-qPCR

Total nucleic acids were extracted from MEF by SDS/Proteinase K treatment at 37 °C followed by phenol-chloroform extraction and ethanol precipitation. Free RNA was removed by RNAse A treatment. DNA was fragmented overnight using HindIII, EcoRI, BsrGI, XbaI, and SspI and pretreated, or not, with RNase H1. For DRIP, R-loops were immunoprecipitated using DNA-RNA hybrids antibody. Bound R-loops were recovered by addition of 50 μl pre-blocked dynabeads protein A magnetic beads (Thermo Fisher Scientific) followed by two washes and elution in an EDTA/SDS-containing buffer. DNA fragments were treated with Proteinase K and recovered with a QIAquick PCR purification kit (Qiagen). Validation of the DRIP was performed by qPCR. Primer pairs used for DRIP analysis are described in Supplemental Information, Table S2.

### DNA immunoprecipitation

DNA immunoprecipitation (DIP) analysis was performed as described earlier (Mohn et al., 2009) with minor adaptations. Briefly, homogenized cells in TE buffer were lyzed overnight in 20 mM Tris–HCl, pH 8.0, 4 mM EDTA, 20 mM NaCl, 1% SDS at 37 °C with 20 µl Proteinase K. Genomic DNA was purified, treated with RNAse A and sonicated with Diagenode Bioruptor to an average DNA length of 300-600 bp. Fragmented DNA was re-purified using Phenol: Chloroform extraction and 4 µg of DNA was immunoprecipitated using antibodies specific against 5mC (Abcam), dsDNA (Abcam) and IgG as a control (Santa Cruz). Precipitated DNA was removed from the beads by incubating with 100 μl of 50 mM Tris–HCl pH 8.0, 10 mM EDTA, 0.5% SDS and Proteinase K for 4 h at 37 °C. DNA was purified using the QIAquick PCR purification kit (Qiagen) and subjected to qPCR. For qPCR, the percentage of input was calculated after subtracting the IgG background.

### Comet-Assay

Comet assay was performed as described by the manufacturer with minor modifications. MEF were treated with doxycycline for 6 h, harvested and mixed with low-melting agarose (Abcam). Mixture was immediately added onto microscopy slides (Abcam). Lysis was performed in alkaline lysis solution (1.2 M NaCl, 100 mM Na_2_EDTA, 0.1% sodium lauryl sarcosinate, 0.26 M NaOH, pH > 13) overnight. Slides were washed and electrophoresed in 0.03 M NaOH, 2 mM Na_2_EDTA (pH ∼12.3) at 1 V/cm for 25 min. DNA was stained with DNA Vista Dye (Abcam) and images were taken with a confocal microscope. Intensities were measured using ImageJ. The tail length and the extended tail moment were calculated as measure for DNA damage.

### Trapping of HMGA2 and dot blot

Dox-inducible MEF were treated with 1 mg/ml dox for 6 h followed by incubation in DPBS at pH 2 for 30 min at 37 °C. Cells were washed with DPBS containing 100 mM NaCNBH_3_ (Sigma), or 100 mM NaCl as a control. Cells were harvested in hypotonic lysis buffer (50 mM Tris-HCl pH 8.0, 2 mM EDTA pH 8.0, 0.1% IGEPAL CA-630 (Sigma), 10% glycerol, 2 mM DTT and protease inhibitor cocktail) and nuclei were resuspended in 50 mM Tris-HCl pH 8.0, 5 mM EDTA pH 8.0, 1% SDS, 2 mM DTT and protease inhibitor cocktail. Nuclei-mix was briefly sonicated on ice and DNA was purified with Phenol: Chloroform. 200 ng of DNA was spotted onto a nitrocellulose membrane (GE Lifescience) for HMGA2 detection.

### GRO-seq, DRIP-seq analysis and R-loop prediction

Reads from GRO-seq and DRIP-seq experiments were downloaded from NCBI, trimmed with trimmomatic and high-quality reads were mapped using Bowtie2 to mouse genome mm10. Genome browser snapshots were created with Homer using makeTagLibrary and makeUCSCfile. Peaks were annotated by using annotePeaks.pl for mm10 from Homer. Correlation of pH2A.X and DNA-RNA hybrids in the top 15% candidates was analyzed using deeptools.

To identify ncRNAs localized in close proximity to the top 15% candidate promoters, an 1 kb window surrounding the TSS was intersected with NONCODEv5_mm10.lncAndGene.bed using bedtools intersect. By using bedtools -S and -s strandness function, antisense and sense orientation was analyzed, respectively.

### DNA nick assay

Genomic DNA of *Hmga2+/+*, *Hmga2-/-* and doxycycline inducible MEF was extracted using the GenElute DNA Miniprep kit (Sigma-Aldrich) according to the protocol provided by the manufacturer. Equal amount of total DNA was applied for Real-time PCR analysis. For detection of DNA nicks, primers for DNA nick assay were designed containing the consensus GT or CT sites specific for DNA nicking enzymes (Gutjahr and Xu, 2014) (Table S2). Nick primers were used with SYBR® Green on the Step One plus Real-time PCR system (Applied Biosystems) and normalized to the Ct values obtained within the surrounding ∼300 bp DNA region amplified with the flanking primers (Table S2). The % of nick DNA was represented as the ratio between: (Nick FWD + Flank RWD) / (Flank FWD + Flank RWD). To determine the directional association of the different ncRNAs associated to the nick DNA area close to the TSS on each target mRNA, we aligned the sequences of the associated ncRNAs using Global Alignment with free end gaps (Geneious 8.1.9, Biomatters Ltd., San Diego, CA). ncRNA:gDNA hybrids were predicted using the RNA hybrid-online server with parameter (MFE<-50 kcal/mol)

### DRIP-WB and DRIP-ChIP

RNA/DNA hybrid IP was performed as described (Cristini et al., 2018) with minor modifications. Non-crosslinked MEF were lysed in 10 mM HEPES pH 7.4, 10 mM KCl, 0.05% NP-40, 1 mM DTT, 25 mM NaF, 0.5 mM Na_3_VO_4_, 40 μg/ml phenylmethylsulfonyl fluoride and protease inhibitor. Nuclei were resuspended in RSB buffer (10 mM Tris-HCl pH 7.5, 200 mM NaCl, 2.5 mM MgCl_2_) with 0.2% sodium deoxycholate, 0.1% SDS, 0.5% Sarkosyl and 0.5% Triton X-100 and sonicated 12 times with the Bioruptor. DNA was measured using Nanodrop and 50 µg of DNA for WB and 25 µg for ChIP was diluted 1:4 in RSB buffer supplemented with 0.5% Trion X-100. Three µg S9.6 antibody was added, and complexes were precipitated using pre-blocked protein A dynabeads (Invitrogen) for two hours while rotating. Beads were washed 4x with RSB supplemented with 0.5% Triton-X100 and eluted in 2x Laemmli buffer for WB or in TE buffer supplemented with 10 mM DTT for ChIP. The standard protocol described above was used for downstream ChIP.

### RNA sequencing and data analysis

Total RNA of *Hmga2+/+* and *Hmga2-/-* MEF treated with water or TGFB1 for 24 h was isolated using Trizol (Invitrogen). RNA was treated with DNase (DNase-Free DNase Set, Qiagen) and repurified using the miRNeasy micro plus Kit (Qiagen). Total RNA and library integrity were verified on LabChip Gx Touch 24 (Perkin Elmer). One µg of total RNA was used as input for SMARTer Stranded Total RNA Sample Prep Kit-HI Mammalian (Clontech). Sequencing was performed on the NextSeq500 instrument (Illumina) using v2 chemistry with 1×75bp single end setup. Raw reads were visualized by FastQC to determine the quality of the sequencing. Trimming was performed using trimmomatic with the following parameters LEADING:3 TRAILING:3 SLIDINGWINDOW:4:15 HEADCROP:5, MINLEN:15. High quality reads were mapped using with HISAT2 v2.1.0 with reads corresponding to the transcript with default parameters. RNA-seq reads were mapped to mouse genome mm10. After mapping, Tag libraries were obtained with MakeTaglibrary from HOMER (default setting). Samples were quantified by using analyzeRepeats.pl with the parameters (mm10-count genes-strand + and –rpkm; reads per kilobase per millions mapped). UCSC known genes with a 1.5-fold change upon TGFB1 treatment in *Hmga2+/+* MEF were classified as TGFB1 inducible and used for downstream analysis. To avoid division through zero, those reads with zero RPKM were set to 0.001.

### ChIP sequencing after Ctrl versus TGFB1 treatment and data analysis

Precipitated DNA samples were purified by QIAquick PCR purification kit (Qiagen) and quantified by Qubit dsDNA HS Assay Kit (Thermo Fisher Scientific). Two ng of DNA was used as input for TruSeq ChIP Library Preparation Kit (Illumina) with following modifications. Instead of gel-based size selection before final PCR step, libraries were size selected by SPRI-bead based approach after final PCR with 18 cycles. In detail, samples were first cleaned up by 1x bead:DNA ratio to eliminate residuals from PCR reaction, followed by 2-sided-bead cleanup step with initially 0.6x bead:DNA ratio to exclude larger fragments. Supernatant was transferred to new tube and incubated with additional beads in 0.2x bead:DNA ratio for eliminating smaller fragments, like adapter and primer dimers. Bound DNA samples were washed with 80% ethanol, dried and resupended in TE buffer. Library integrity was verified with LabChip Gx Touch 24 (Perkin Elmer). Sequencing was performed on the NextSeq500 instrument (Illumina) using v2 chemistry with 1×75bp single end setup. Raw reads were visualized by FastQC to determine the quality of the sequencing. Trimming was performed using trimmomatic with the following parameters LEADING:3 TRAILING:3 SLIDINGWINDOW:4:15 HEADCROP:4, MINLEN:15.

High quality reads were mapped by using with bowtie2 (Langmead and Salzberg, 2012) to mouse genome mm10. For downstream analysis, reads were scaled based on their read counts and normalized by subtracting reads of the corresponding inputs using deeptools.

### Crossing of murine TGFB1 with human IPF data and GSEA

IPF RNA-seq samples from GSE116086 were remapped by the help of bowtie2 to human genome version hg38. Differential gene expression was analyzed using DEseq2 (default) (doi: 10.1186/s13059-014-0550-8.). Human gene name was converted to mouse (mgi_symbol) by the use of getLDS from biomaRt program. IFP-RNA-seq was crossed by mgi_symbol with the Hmga2-RNA-seq after TGFB1 treatment in *Hmga2+/+* MEF. The log2 fold change (log2FC) both RNA-seq was used to perform a 2D kernel density plot by the help of the function kde2d from MASS package v7.3-51.4 with the number of grip points 50. Gene enrichment set analysis (GSEA) was obtained using fgsea (parameters minSize = 10, nperm=1000) taken the “h.all.v7.0.symbols.gmt” as pathway database. PlotEnrichment was used to plot the two most enriched pathways from the Up-regulated genes from either IPF-RNA-seq or Hmga2-RNA-seq after TGFB1 treatment.

### Migration and proliferation assays

Lung fibroblasts, to be assessed for cellular proliferation, were cultured either in 96-well or 48-well plates. Fibroblast proliferation was determined using colorimetric BrdU incorporation assay kit (Roche) according to manufacturer’s instructions. Absorbance was measured at 370 nm with reference at 492 nm in a plate reader (TECAN). Depending on the experiment, proliferation of cells was plotted either as the difference of absorbance at 370 and 492 nm (A370 nm–A492 nm) or as a percentage of absorbance compared to control cells absorbance.

### Collagen assays

Total collagen content was determined using the Sircol Collagen Assay kit (Biocolor, Belfast, Northern Ireland). Equal amounts of protein lysates from Ctrl and IPF human lung fibroblasts were added to 1 ml of Sircol dye reagent, followed by 30 min of mixing. After centrifugation at 10,000 × g for 10 min, the supernatant was carefully aspirated, and 1 ml of Alkali reagent was added. Samples and collagen standards were then read at 540 nm on a spectrophotometer (Bio-Rad). Collagen concentrations were calculated using a standard curve generated by using acid-soluble type 1 collagen.

### Hydroxyproline measurements

Hydroxyproline levels in human lung fibroblasts were determined using the QuickZyme Hydroxyproline Assay kit (Quickzyme Biosciences). The cells and lung tissue were separately homogenized in 1 ml 6 N HCl with a Precellys tissue homogenizer (2 × 20 s, 3,800 g). The homogenate was then hydrolyzed at 90 °C for 24 h. After centrifugation at 13,000 g for 10 min, 100 μl from the supernatant was taken and diluted 1:2 with 4 N HCl. 35 μl of this working dilution was transferred to a 96-well plate. Likewise, a hydroxyproline standard (12.5–300 μM) was prepared in 4 N HCl and transferred to the microtiterplate. Following addition of 75 μl of a chloramine T-containing assay buffer, samples were oxidized for 20 min at room temperature. The detection reagent containing p-dimethylaminobenzaldehyde was prepared according to the manufacturer’s instruction and 75 μl added to the wells. After incubation at 60 °C for 1 h, the absorbance was read at 570 nm with a microtiter plate reader (Infinite M200 Pro, Tecan) and the hydroxyproline concentration in the sample was calculated from the standard curve and related to the employed amount of lung tissue. The hydroxyproline content in lung tissue is given as μg hydroxyproline per mg lung tissue.

### Experiments with human PCLS

PCLS were prepared from tumor-free lung explants from patients who underwent lung resection for cancer at KRH Hospital Siloah-Oststadt-Heidehaus or the Hanover Medical School (both Hanover, Germany). Tissue was processed immediately within 1 day of resection as described before (Wujak et al., 2017). Briefly, human lung lobes were cannulated with a flexible catheter and the selected lung segments were inflated with warm (37 °C) low-melting agarose (1.5%) dissolved in DMEM Nutrient Mixture F-12 Ham supplemented with l-glutamine, 15 mM HEPES without phenol red, pH 7.2–7.4 (Sigma Aldrich), 100 U/ml penicillin, and 100 µg/mL streptomycin (both from Biochrom). After polymerization of the agarose solution on ice, tissue cores of a diameter of 8 mm were prepared using a sharp rotating metal tube. Subsequently, the cores were sliced into 300–350 µm thin slices in DMEM using a Krumdieck tissue slicer (Alabama Research and Development). PCLS were washed 3× for 30 min in DMEM and used for experiments. Viability of the tissue was assessed by an LDH Cytotoxicity Detection Kit (Roche) according to the manufacturer’s instruction. For IPF resolution experiments, human IPF PCLS were treated with 50 or 100 μM FACTin for 72 h and the medium with FACTin replenished every 24 h.

### Immunofluorescence staining in PCLS

PCLS from IPF patients were fixed with acetone/methanol (Roth) 50:50 by volume for 20 min, blocked for 1 h with 5% bovine serum albumin (w/v, Sigma) in 1x PBS, pH 7.4. Cells were then incubated with primary antibody overnight at 4 °C. After incubation with a secondary antibody for 1h, nuclei were DAPI stained and PCLS were examined with a confocal microscope (Zeiss). Antibodies used were specific for DNA-RNA hybrid (S9.6, Kerafast), COL1A1 (Sigma), ACTA2 (Sigma), FN1 (Milippore), VIM (Cell Signaling), pH2A.X (Millipore) and HMGA2 (SantaCruz). Alexa 488, Alexa555 or Alexa 594 tagged secondary antibodies (Invitrogen) were used. DAPI (Sigma Aldrich) used as nuclear dye.

### Publicly available datasets

We used a number of publicly available datasets to aid analysis of our data: pH2A.X and HMGA2 ChIP-Seq data: GEO: GSE63861 (Singh et al., 2015) DRIP-Seq in NIH/3T3 and E14 cells data: GEO: GSE70189 (Sanz et al., 2016) Nuclear RNA-seq in Ctrl and human IPF hLF: GEO: GSE116086 (Rubio et al., 2019) GRO-seq in Mouse embryonic fibroblasts: GEO: GSE76303 (Busslinger et al., 2017)

### Quantification and Statistical Analysis

A summary of the *P* values and the statistical tests used in the different experiments can be found in Table S3. Further details of statistical analysis in different experiments are included in the Figures and Figure legends. Briefly, protein enrichment on chromatin and expression analysis of samples were analyzed by next generation sequencing. Tree independent experiments of the mass spectrometry-based proteomic approach were performed. For the rest of the experiments presented here, samples were analyzed at least in triplicates and experiments were performed three times. Statistical analysis was performed using GraphPad Prism 5 and Microsoft Excel. Data in bar plots are represented as mean ± standard error (mean ± s.e.m.). T-tests were used to determine the levels of difference between the groups and *P* values for significance. *P* values after one- or two-tailed t-test, \**P* ≤ 0.05; \*\**P* ≤ 0.01 and \*\*\**P* ≤ 0.001. In the box plots of Figures 1B, 2D, 3F, 6B and S6B, *P* values were determined using Wilcoxon-Mann-Whitney test. In the 2D Kernel Density plot presented in Figure 7A the statistical significance was calculated using DESeq2’s integrated Wald test.

### Data and Software Availability

Supplemental Information is linked to the online version of the paper at http://www.cell.com. In addition, sequencing data of ChIP, RNA and MNase have been deposited in NCBI’s Gene Expression Omnibus (Edgar et al., 2002) and is accessible through GEO Series with accession number GSE141272. To access the private session, reviewer can use the Token: wrkxkkqefhwnfon. The mass spectrometry-based interactome data have been deposited into the PRIDE (https://www.ebi.ac.uk/pride/archive/) archive and assigned to the project accession PXD016586. Reviewer can access the data using Username: and Password: sMtpjWKM.

## SUPPLEMENTAL INFORMATION

**Figure S1:**
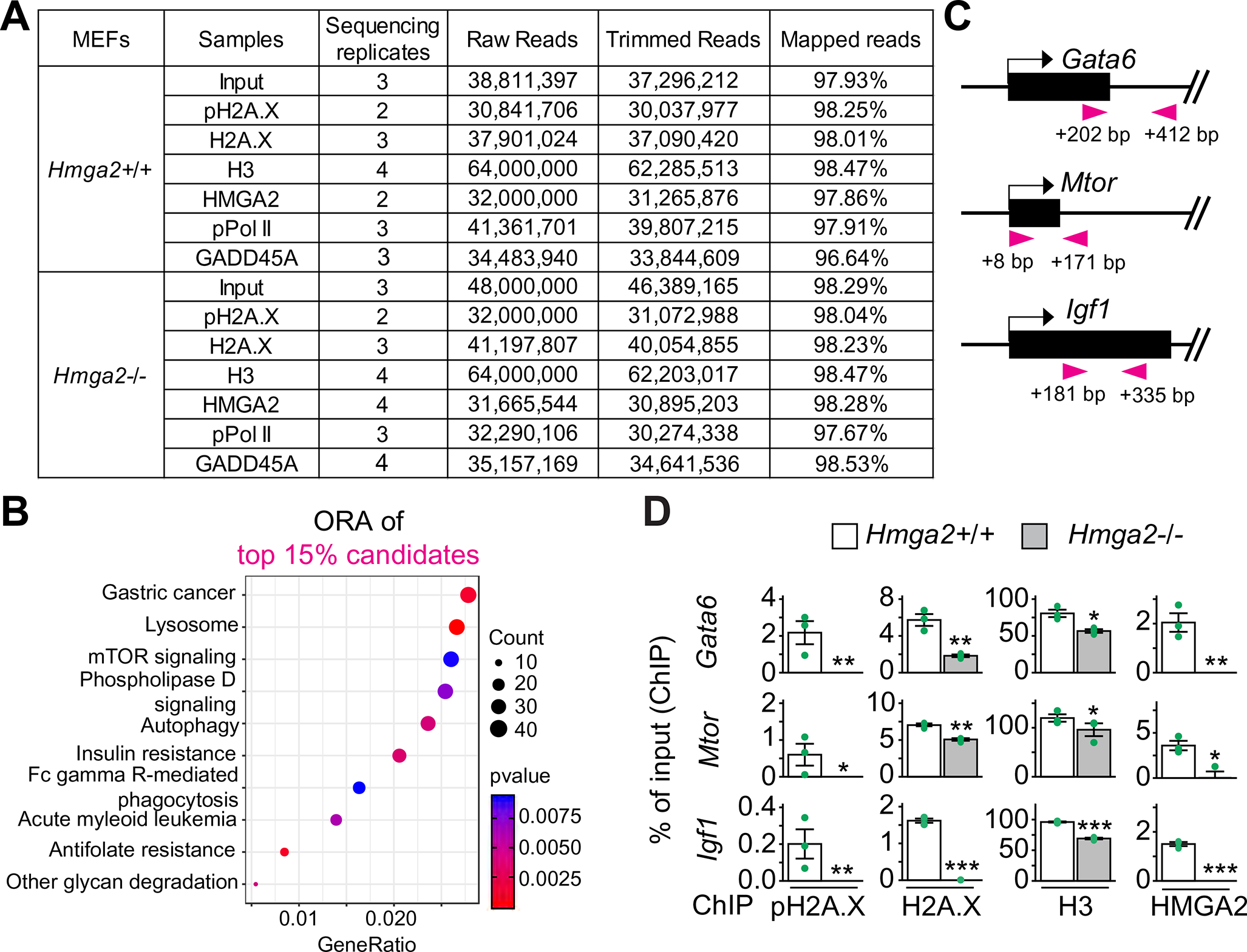
HMGA2 is required for pH2A.X deposition at TSS. Related to Figure 1. (A) Description of the ChIP-seq data set supports the quality of the experiment. (B) KEGG-based enrichment analysis of top 15 % candidate genes using clusterProfiler. (C) Schematic representation of the 5’ genomic region of *Igf1*, *Gata6* and *Mtor* showing exons (black boxes), introns (lines), arrows, direction of the genes and location of primer pairs (arrowheads) used for ChIP analysis. Number represents distance of the 5’ primer to the TSS. (D) ChIP analysis of *Gata6*, *Mtor* and *Igf1* TSS using specific antibodies as indicated and chromatin isolated from *Hmga2+/+* and *Hmga2-/-* MEF. In all plots, data are displayed as means ± s.e.m (n = 3 independent experiments); asterisks, *P* values after *t-*Test, \*\*\**P* ≤ 0.001; \*\**P* ≤0.01; \**P* ≤ 0.05; ns, non-significant. The statistical test values of each plot are shown in Supplementary Table S3.

**Figure S2:**
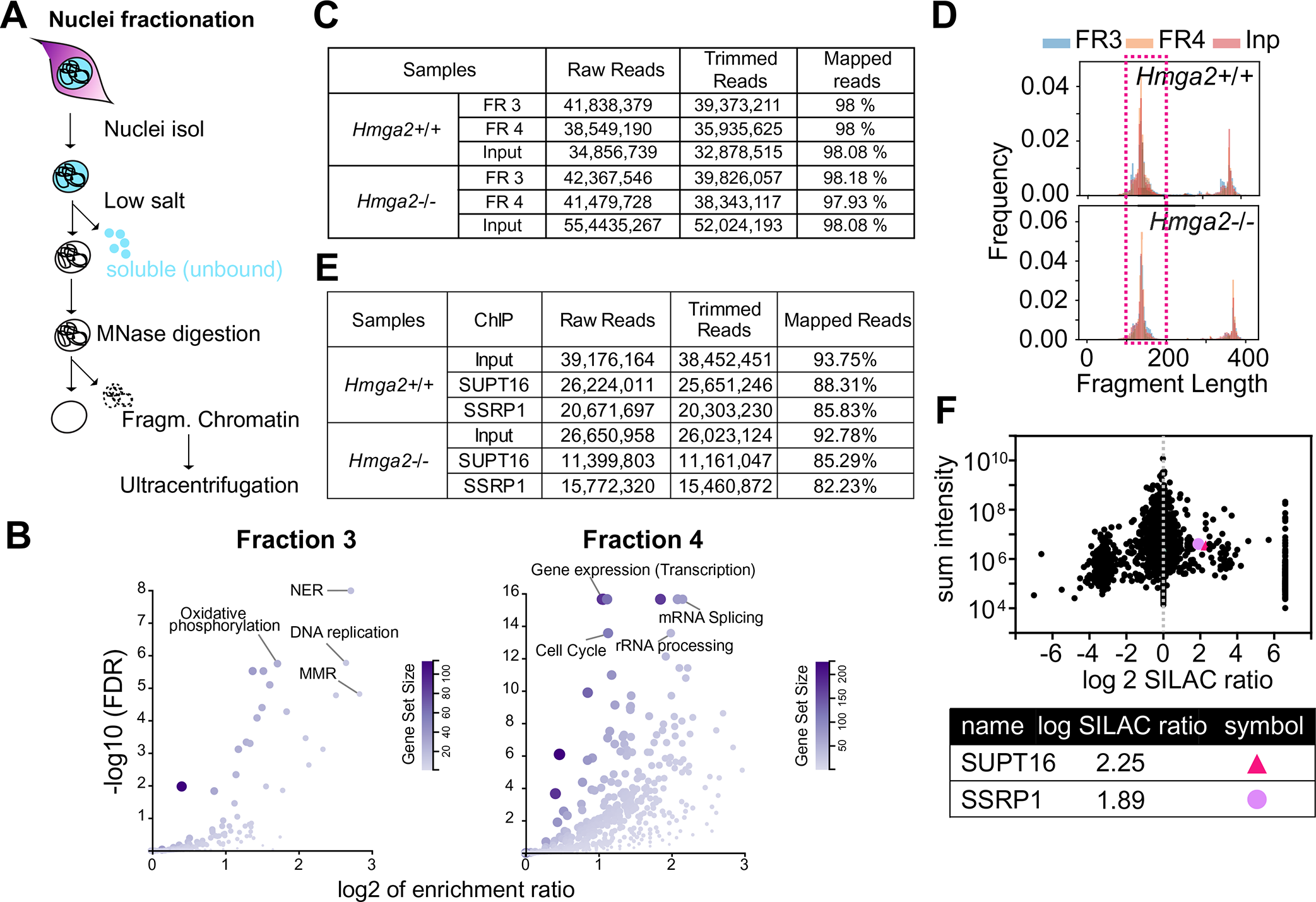
Hmga2 KO decreases binding of proteins to chromatin. Related to Figure 3. (A) Schematic representation of the experimental outline. Chromatin was prepared and fractionated by micrococcus nuclease digestion. Samples were loaded onto a sucrose gradient and were separated by ultracentrifugation. (B) Left, KEGG-based and, right, Reactome-based enrichment analysis of proteins showing significant enrichment in *Hmga2+/+* as compared to *Hmga2-/-* MEF in fraction 3 and 4 using WEB-based gene set analysis toolkit, respectively. (C, E) Description of the MNase-seq (C), SUPT16 and SSRP1 ChIP-seq (E) data set supports the quality of the experiments. (D) Frequency of fragment length distribution of reads obtained from fraction 3 and 4 after paired-end sequencing. Reads with a length of 100 to 200 bp were selected for further analysis. (F) HMGA2-interacting proteins identified by mass spectrometry analysis published by (Singh et al., 2015). Scatter plot between SILAC ratios and peak intensities (top); selected proteins with corresponding log2 SILAC ratios of SUPT16 and SSRP1 (bottom).

**Figure S3:**
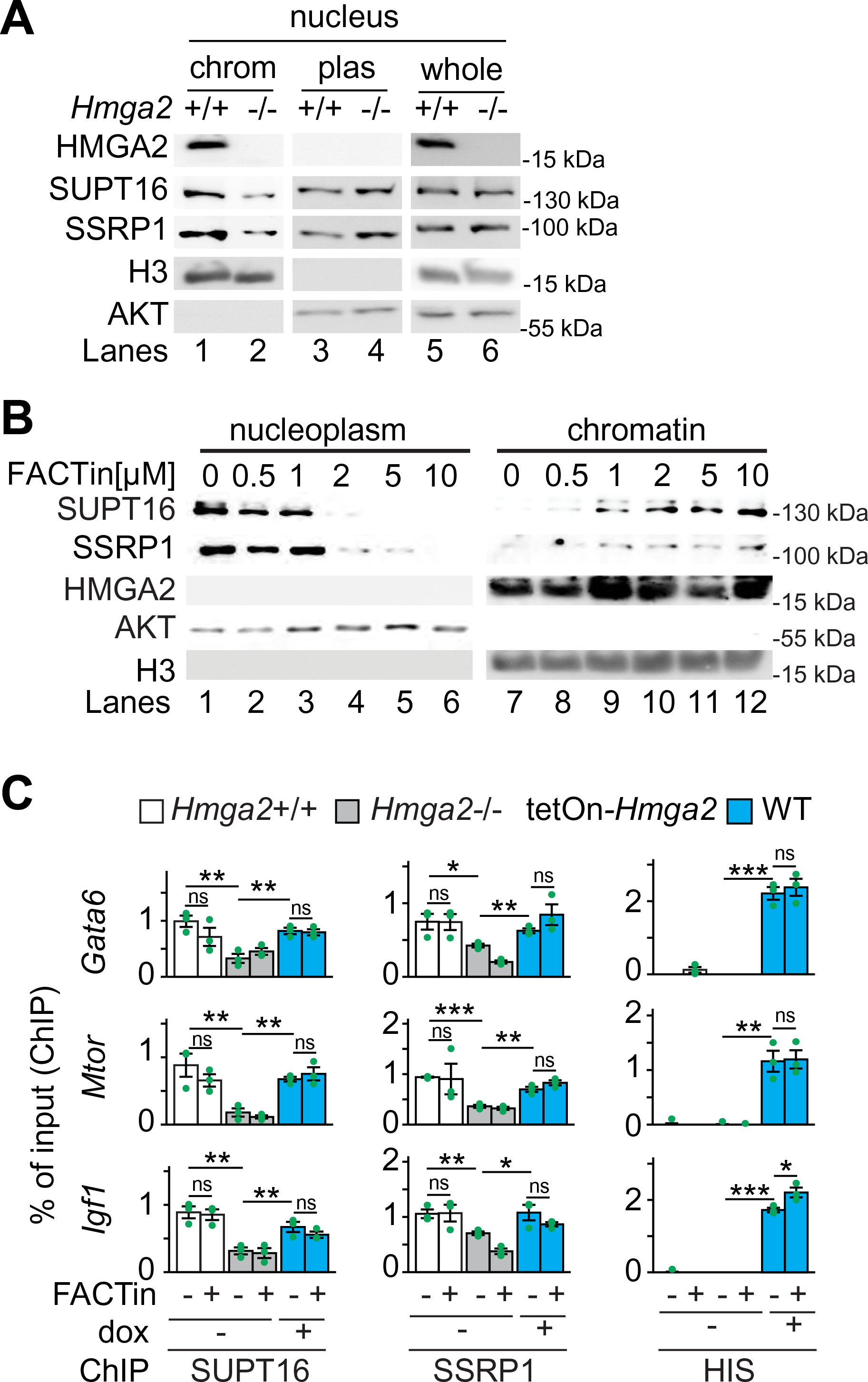
HMGA2 and FACTin increases FACT complex binding to chromatin. Related to Figure 4. (A) WB analysis of chromatin (chr), nucleoplasm (nuc pl) and nuclear lysates (nuc lys) from *Hmga2+/+* and *Hmga2-/-* MEF using the indicated antibodies. H3 and AKT were used as markers for chromatin and nucleoplasm, respectively. (B) *Hmga2+/+* MEF were treated for 2 h with the indicated concentrations of CBLC000 trifluoroacetate (FACTin). Afterwards, nucleoplasm and chromatin were isolated and analyzed by WB using the noted antibodies. AKT and H3 were used as nucleoplasm and chromatin markers, respectively. (C) ChIP-based promoter analysis of *Gata6*, *Mtor* and *Igf1* using the indicated antibodies and chromatin from *Hmag2+/+*, *Hmag2-/-* MEF, as well as *Hmga2-/-* MEF that were stably transfected with a tetracycline-inducible expression construct (tetOn) for WT *Hmga2-myc-his*. MEF were treated with doxycycline and FACT inhibitor (FACTin; CBLC000 trifluoroacetate) as indicated. Data are displayed as means ± s.e.m (n = 3 independent experiments); asterisks, *P* values after *t*-Test, \*\*\**P* ≤ 0.001; \*\**P* ≤ 0.01; \**P* ≤ 0.05; ns, non-significant. The statistical test values of each plot are shown in Supplementary Table S3.

**Figure S4:**
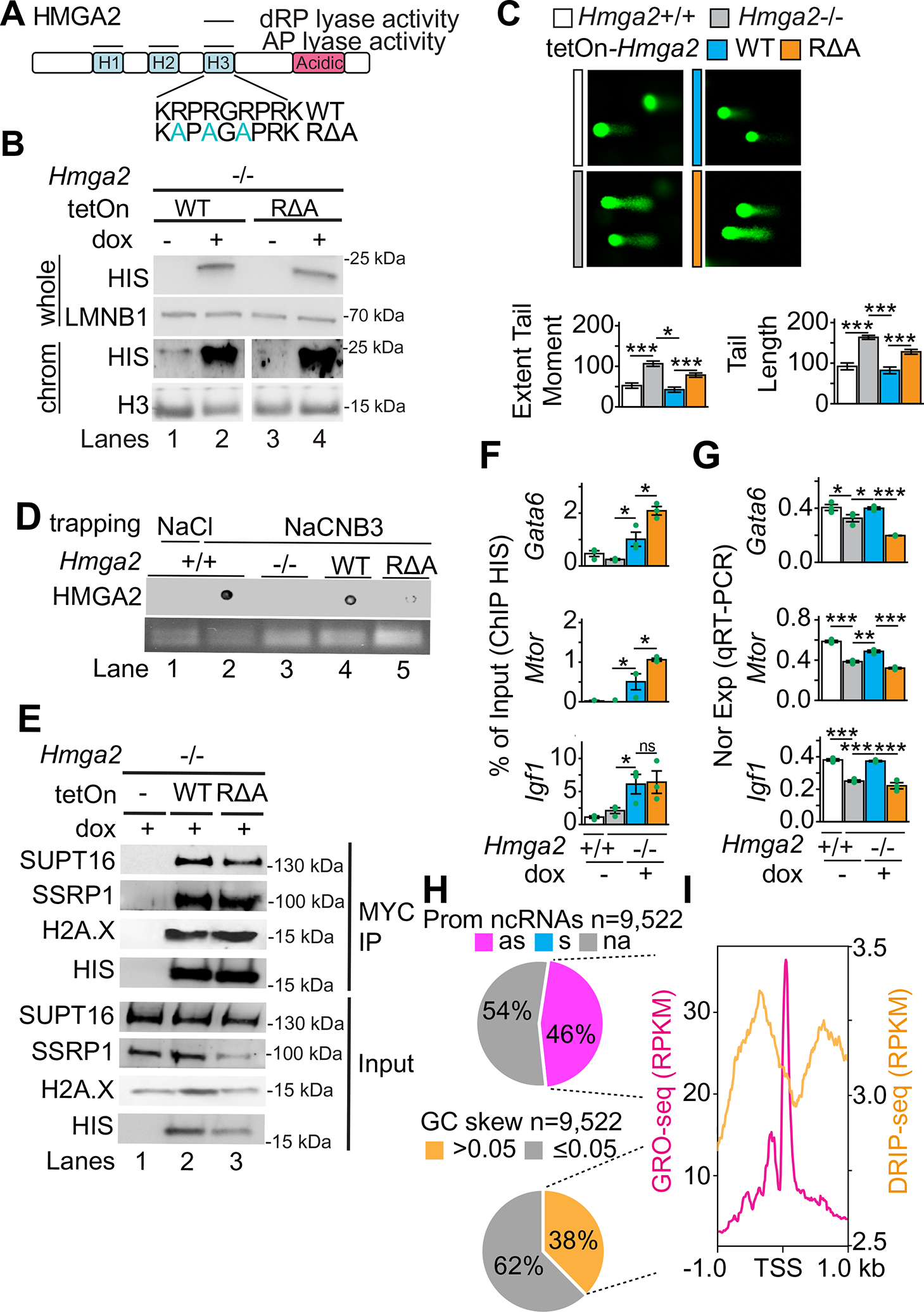
Lyase activity-dependent pH2A.X deposition is required for pPol II enrichment and gene transcription. Related to Figure 4. (A) Schematic representation of HMGA2 comprising AP lyase activity within the three AT-hook domains (H1-3) and a 5’-desoxyribose phosphate (dRP) lyase activity. Arginine (R) residues were mutated to alanines to abolish lyase activity in the third hook domain (RΔA mutant). (B) WB analysis of nuclear lysate (top) and chromatin (bottom) of *Hmga2-/-* MEF that were stably transfected with a tetracycline-inducible expression construct (tetOn) either for WT *Hmga2-myc-his* or the lyase-deficient mutant RΔA *Hmga2-myc-his* using the indicated antibodies. LMNB1 and H3 were used as a loading control. (C) Top, representative images of comet assay using *Hmga2+/+*, *Hmga2-/-* MEF, as well as *Hmga2-/-* MEF that were stably transfected with a tetracycline-inducible expression construct (tetOn) either for WT *Hmga2-myc-his* or the lyase-deficient mutant RΔA *Hmga2-myc-his*. Cells were treated for 6 h with doxycycline. Bottom, quantification of extent tail moment and tail length in imaged MEF. (D) Dot-blot confirming the impaired lyase activity of the RΔA mutant. A control for the amount of DNA loaded is shown below the blot. (E) Immunoprecipitation of HMGA2 from nuclear protein extracts of *Hmga2-/-* MEF, as well as *Hmga2-/-* MEF that were stably transfected with a tetracycline-inducible expression construct (tetOn) either for WT *Hmga2-myc-his* or the lyase-deficient mutant RΔA *Hmga2-myc-his* using MYC-coated magnetic beads. Co-immunoprecipitated proteins were analyzed by WB using the indicated antibodies. Input (Inp), 4% of material used for the IP. (F) ChIP analysis of HMGA2 targets using a HIS-tag specific antibody with chromatin isolated from *Hmga2+/+*, *Hmga2-/-* MEF, as well as *Hmga2-/-* MEF that were stably transfected with a tetracycline-inducible expression construct (tetOn) either for WT *Hmga2-myc-his* or the lyase-deficient mutant RΔA *Hmga2-myc-his*. Cells were treated for 4h with doxycycline as indicated. (G) QRT-PCR analysis using *Hmga2+/+*, *Hmga2-/-* MEF, as well as *Hmga2-/-* MEF that were stably transfected with a tetracycline-inducible expression construct (tetOn) either for WT *Hmga2-myc-his* or the lyase-deficient mutant RΔA *Hmga2-myc-his*. Cells were treated for 6h with doxycycline. In all plots, data are displayed as means ± s.e.m (n = 3 independent experiments); asterisks, *P* values after *t-*Test, \*\*\**P* ≤ 0.001; \*\**P* < 0.01; \**P* ≤ 0.05; ns, non-significant. The statistical test values of each plot are shown in Supplementary Table S3. (H) Top, pie chart representing the promoter associated ncRNAs of the top 15 % candidate genes in antisense and sense orientation or do not contain annotated ncRNAs. As, antisense; s, sense; na, not-annotated. Bottom, pie chart highlighting the distribution of the top 15 % genes with a high GC skew (>0.05). (I) Aggregate plot representing the global distribution of nascent RNA (GRO-seq) in WT MEF of the top 15 % genes with associated antisense ncRNA and RNA-DNA hybrids (DRIP-seq) in NIH/3T3 murine fibroblasts with high GC skew. Reads were normalized using an RPKM measure. TSS, transcription start site; kb, kilobases).

**Figure S5:**
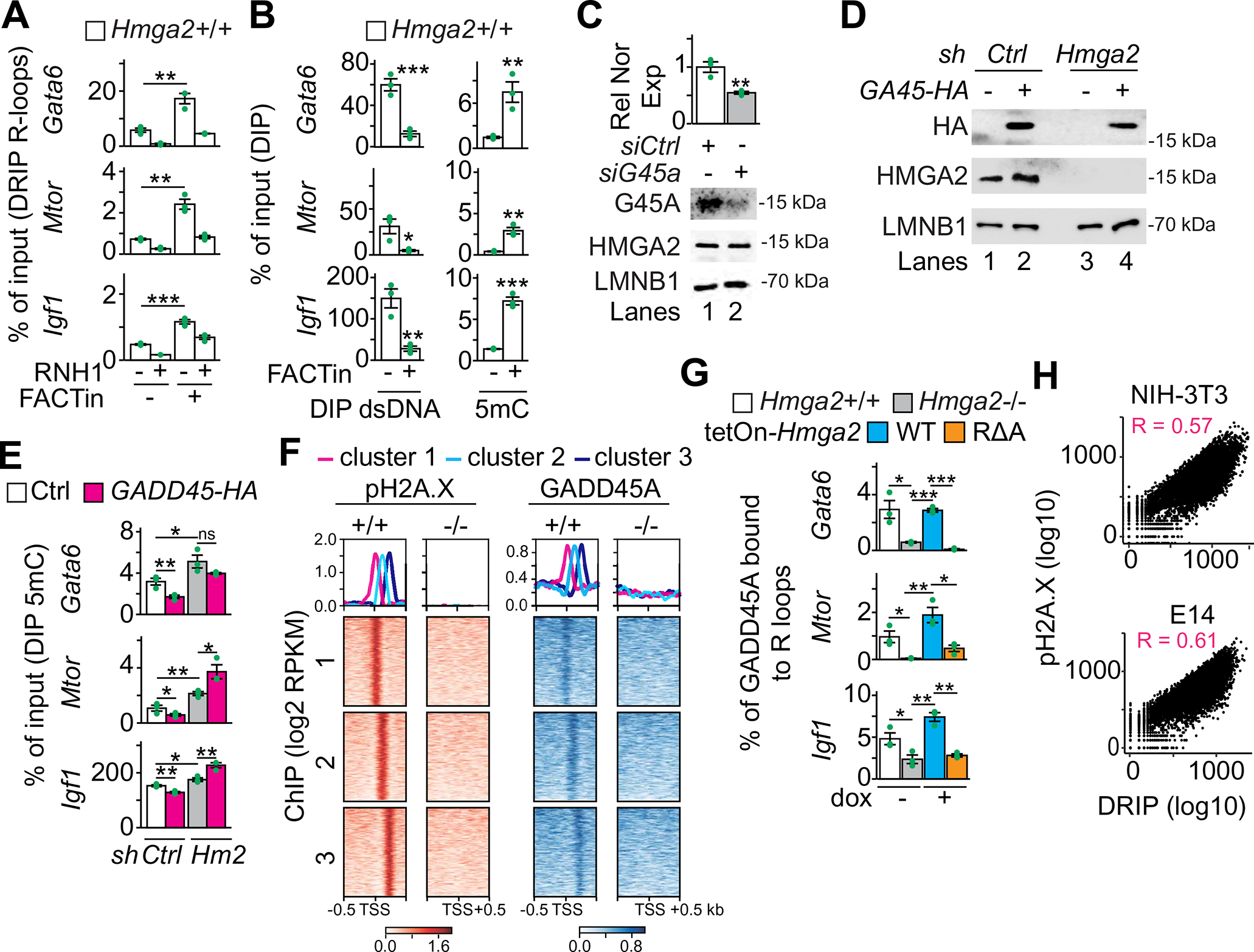
R-Loops correlate with pH2A.X enrichment and GADD45A-induced DNA demethylation. Related to Figure 5. (A-B) Analysis of R-loops (A), dsDNA and 5mC (B) in the TSS of HMGA2 targets in *Hmga2+/+* treated for 4 h with FACTin as indicated. (C) QRT-PCR analysis and WB for *Gadd45a* expression in *Hmga2+/+* MEF after siRNA-mediated KD. (D, E) WB analysis (D) and DIP for 5mC (E) after overexpression of *GADD45*A tagged with HA in MLE-12 cells that were stably transfected either with a control (scramble, *scr*) or an *Hmga2*-specific short hairpin DNA (*sh*) construct. (F) Aggregate plot and heat maps for pH2A.X and GADD45A enrichment at the TSS ± 0.5kb of the top 15 % candidates in *Hmga2+/+* and *Hmga2-/-* MEF within the same clusters identified in Figure 2E. (G) DRIP followed by sequential ChIP for GADD45A in *Hmga2+/+*, *Hmga2-/-* MEF, as well as *Hmga2-/-* MEF that were stably transfected with a tetracycline-inducible expression construct (tetOn) either for WT *Hmga2-myc-his* or the lyase-deficient mutant RΔA *Hmga2-myc-his*. Cells were treated for 4h with doxycycline. In all plots, data are displayed as means ± s.e.m (n = 3 independent experiments); asterisks, *P* values after *t*-Test, \*\*\**P* ≤ 0.001; \*\**P* ≤ 0.01; \**P* ≤ 0.05; ns, non-significant. (H) Correlation analysis of pH2AX mapped reads with DNA-RNA hybrids (mapped reads; DRIP-seq) in NIH/3T3 and E14 cells using the top 15 % genes. R, Pearson correlation. The statistical test values of each plot are shown in Supplementary Table S3.

**Figure S6:**
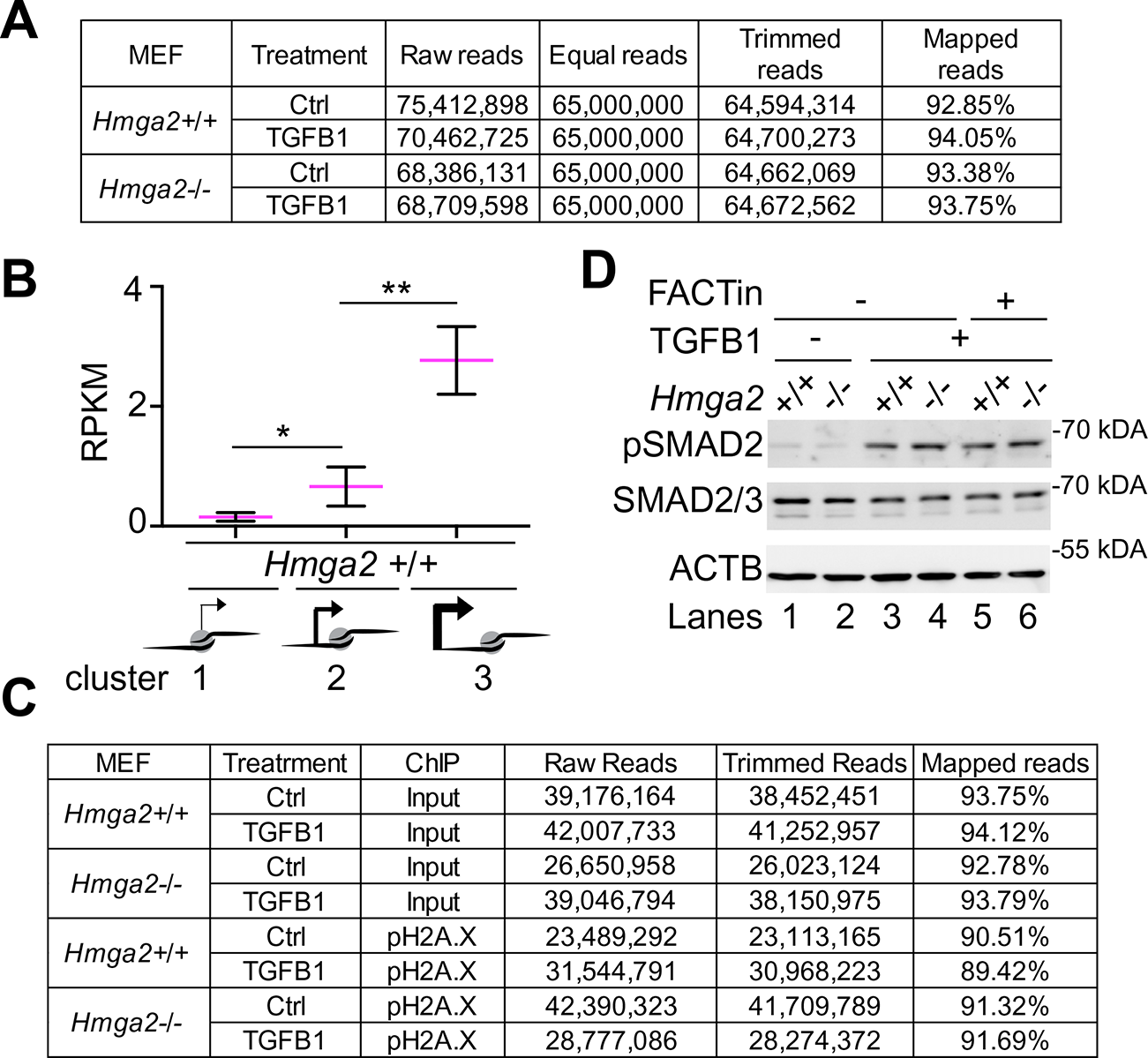
GADD45A is required for TGFB1-enhanced active DNA methylation. Related to Figure 6. (A, C) Descriptions of the RNA-seq data and TGFB1 ChIP-seq set support the quality of the experiments. (B) TGFB1-inducible genes in WT MEF were selected from Figure 2F and their expression was analyzed in *Hmga2+/+* and *Hmga2-/-* MEF measured by RPKM. *P* values after Wilcoxon-Mann-Whitney test, ** *P* ≤ 0.01; * *P* ≤ 0.05. The statistical test values of each plot are shown in Supplementary Table S3. (D) WB analysis of phosphorylated Smad2 (Ser465/467), pSMAD) and total SMAD2/3 of *Hmga2+/+* and *Hmga2-/-* MEF treated with TGFB1 alone or in combination with FACTin.

**Figure S7:**
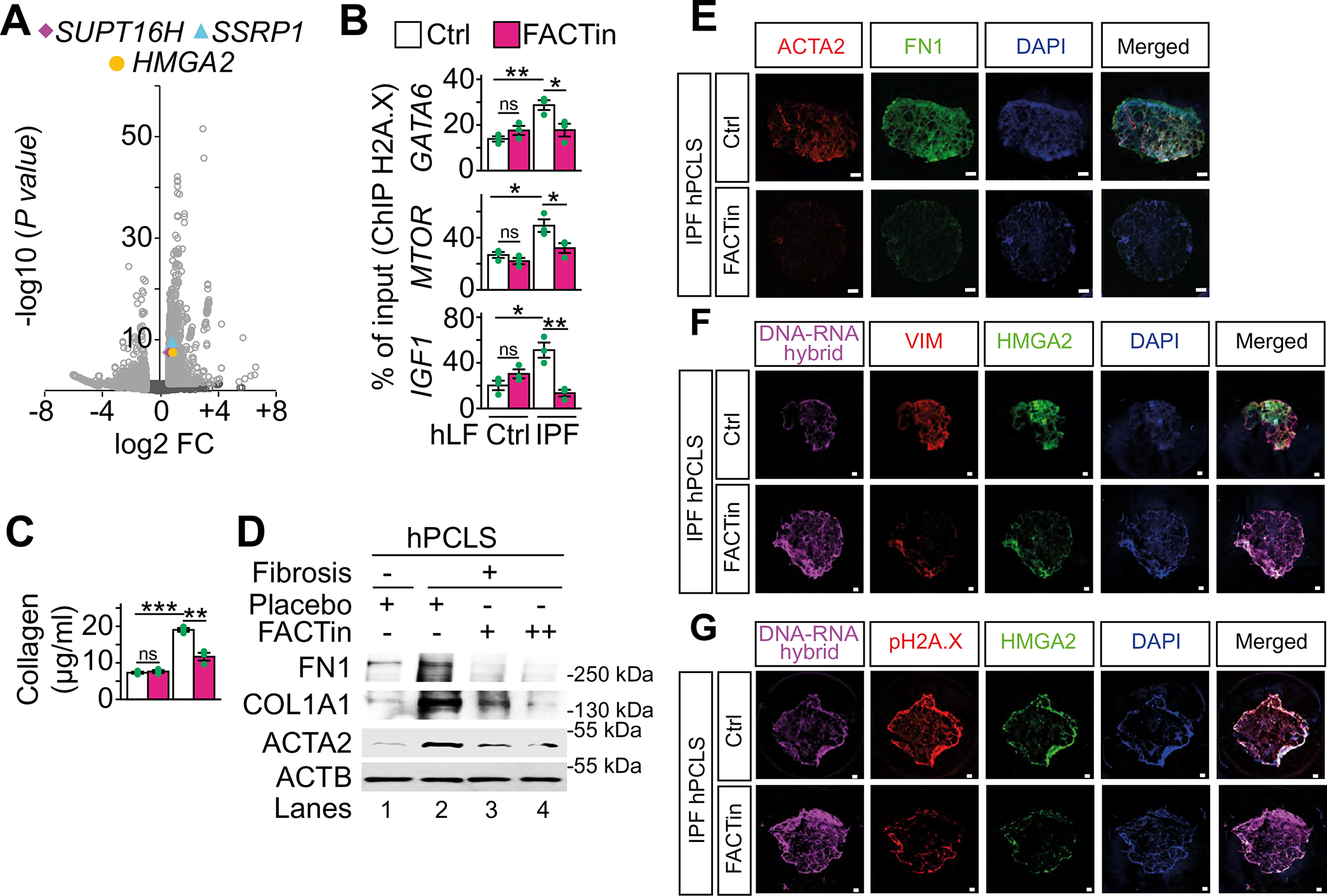
FACTin reduces fibrotic markers and pH2A.X levels in PCLS. Related to Figure 7. (A) Volcano plot representing the significance (−log10 *P* values after Wald t-test) vs. expression fold change (log2 expression ratio) between two Ctrl and two IPF patients. Fold changes of *SUPT16H*, *SSRP1* and *HMGA2* in IPF are highlighted. Light grey color marks genes with a FC>1.5 and *P*<0.05. (B) ChIP-based promoter analysis of selected HMGA2 target genes using H2A.X-specific antibody and chromatin from hLF treated as in Figure 7D. (C) Quantification of collagen content in Ctrl and IPF hLF treated with Ctrl or FACTin for 24h. In all bar plots, data are shown as means ± s.e.m. (*n* = 3 independent experiments); asterisks, *P* values after *t*-Test, *** *P* ≤ 0.001; ** *P* ≤ 0.01; * *P* ≤ 0.05; ns, non-significant. The statistical test values of each plot are shown in Supplementary Table S3. (D) WB analysis of IPF markers in Ctrl and IPF human precision-cut lung slices (hPCLS) treated with FACTin using the indicated antibodies. (E-G) Representative pictures of confocal microscopy after immunostaining using ACTA2, FN1, S9.6, VIM, HMGA2 or pH2A.X-specific antibody in hPCLS from IPF patients. The hPCLS were treated as in Figure 7I. ACTA2, smooth muscle actin alpha 2; FN1, fibronectin; VIM, vimentin; DAPI, nucleus. Scale bars, 500 μm.

## TABLES

Supplementary Table 1 (S1): Data sheets containing ChIP-seq, RNA-seq and mass spectrometry based proteomic presented in Figures 1C, 2E, 2F, 3B, 6A and 7A.

Supplementary Table 2 (S2): Primer sequences and sequences of shDNA constructs.

Supplementary Table 3 (S3): Statistical summary containing the values for statistical significance and the implemented statistical tests in all plots presented in the article.

